# Survival of the “unfittest”: clinical emergence of hyper-multidrug-resistant *Nakaseomyces glabratus* with rare nonfunctional Erg3 and Erg11 and severely impaired fitness

**DOI:** 10.1101/2025.02.05.636719

**Authors:** Hans Carolus, Vladislav Biriukov, Jolien Vreys, Celia Lobo Romero, Juan Paulo Herrera Avila, Rudy Vergauwen, Dimitrios Sofras, Giel Vanreppelen, Lore Vinken, Basil Britto Xavier, Toni Gabaldón, Katrien Lagrou, Reinout Naesens, Patrick Van Dijck

## Abstract

**Background:** *Nakaseomyces glabratus* (*Candida gabrata*) poses a significant clinical challenge due to common drug resistance. We report a case of a complicated urinary tract infection (UTI) progressing to prostatitis and urosepsis, with the emergence of a hyper-multidrug-resistant isolate with low stress tolerance, slow growth and a short life span. This study elucidates the genetic mechanisms and phenotypic characteristics underlying antifungal hyper-resistance with strong fitness trade-offs, and explores potential alternative therapies for resistant UTI’s.

**Methods:** Whole-genome sequencing was performed to identify resistance-associated mutations and gene knock-out strains were generated to assess the relative impact of putative loss-of-function (LoF) mutations on antifungal resistance, fitness and membrane sterol composition. Drug susceptibility testing of the antibiotic nitroxoline and related compounds was conducted to evaluate it as a therapeutic alternative and study the mechanism of action.

**Findings:** Loss-of-function mutations in *ERG3* and *ERG11* were identified and linked to the accumulation of 4,14-dimethylzymosterol and lanosterol instead of ergosterol. Engineered *ERG3Δ+ERG11Δ* strains recapitulated the clinical isolate’s hyper-multidrug resistance and associated fitness deficits. While *ERG3Δ* strains showed no resistance but enhanced thermotolerance, *ERG11Δ* and *ERG3Δ+ERG11Δ* strains exhibited multidrug resistance with severe fitness trade-offs. Interestingly, *ERG3Δ+ERG11Δ* strains showed mild resistance to flucytosine, but an additional *FUR1* mutation in the clinical isolate most probably underlies hyper-resistance to flucytosine. The UTI antibiotic nitroxoline demonstrated high antifungal activity against all strains, and the LoF of *ERG3* and/or *ERG11* induced collateral sensitivity to this drug. Testing of related compounds suggest a mode of action beyond iron chelation.

**Interpretation:** This case demonstrates that hyper-resistant strains of *N. glabratus* can emerge despite significant fitness costs and persist under prolonged antifungal therapy in specific clinical settings. These findings underscore the importance of vigilant antifungal resistance monitoring and highlight nitroxoline as a promising alternative treatment for complicated fungal UTIs. These results challenge the notion that strains with fitness deficits are clinically irrelevant and emphasize the need for novel therapeutic strategies including repurposed agents.

## Introduction

Fungal infections form a major global health burden, causing around 6.5 million invasive infections and 3.8 million deaths annually. *Candida* species stand out among fungal pathogens, due to their significant morbidity and mortality rates and increasing resistance to the limited arsenal of antifungal drugs. They account for an estimated 1 565 000 cases of bloodstream infections and invasive candidiasis, resulting in 995 000 associated deaths each year, reflecting a mortality rate of 63.6% (1). The spectrum of species causing candidemia has shifted from the traditionally drug-susceptible *Candida albicans* to non-albicans *Candida* (NAC) species, such as *Candida glabrata, Candida tropicalis, Candida parapsilosis* and *Candida auris* (2)*. C. glabrata*, which was recently renamed *Nakaseomyces glabratus* (3), causes 20 to 25% of invasive *Candida* infections in Western Europe and the USA (2) and is now one of the leading causes of candidemia in many healthcare settings, approaching the prevalence of *C. albicans*, which historically has been the dominant species (3).

In nosocomial settings, *N. glabratus* is implicated in approximately 21% of *Candida* urinary tract infections (UTIs) cases, second only to *C. albicans*, which causes approximately 49% of cases (4). While asymptomatic candiduria is often benign, symptomatic cases can manifest as cystitis, pyelonephritis, or other complications, particularly in high-risk patients. Fluconazole is the standard first-line treatment, but effectiveness of this drug against *N. glabratus* is limited due to intrinsic resistance to azoles (4). Due to the poor urinary concentrations achieved by echinocandins, amphotericin B deoxycholate or flucytosine, are recommended for UTIs of fluconazole-resistant NAC species (4–6).

Although amphotericin B has been a mainstay in antifungal therapy for over 70 years, resistance to this drug remains rare, which might be partially explained by the fitness trade-offs associated with the mechanisms of amphotericin B resistance (7–10). One elusive mechanism of hyper-resistance is the loss of function (LoF) of *ERG11,* which leads to the accumulation of toxic 14-methyl-ergosta-8,24(28)-diene-3,6-diol. The epistatic LoF of *ERG3* can mitigate this toxicity. Therefore, the combined LoF of both *ERG11* and *ERG3* has been identified as a putative mechanism of high amphotericin B resistance in several *in vitro* experimental evolution (7–9) and gene knock-out studies (11, 12). Nevertheless, this mechanism of resistance has been linked to severe fitness trade-offs (7, 9) which have been predicted to render the pathogen avirulent (8, 9). While LoF mutations in both *ERG3* and *ERG11* have been documented in clinical isolates of *C. tropicalis* (13), this genotype has not been reported in other *Candida* species. Here, we describe the case of a patient that developed a complicated *N. glabratus* UTI progressing to prostatitis and urosepsis. Following treatment with fluconazole, liposomal amphotericin B, amphotericin B deoxycholate, flucytosine and anidulafungin, the emergence of a hyper-multidrug-resistant isolate with LoF mutations in *ERG3* and *ERG11* led to significant therapeutic challenges. This study investigates the genetic basis of amphotericin B resistance, its associated fitness trade-offs, and the potential for alternative therapies, providing insights into antifungal resistance mechanisms and clinical management strategies.

## Results

### Clinical case

An 80-year-old male with a history of type 2 diabetes and stable ischemic cardiopathy presented to the ZAS urology clinic with several weeks of lower urinary tract symptoms and persistent isolation of *N. glabratus* in urine cultures (**Figure 1A**). The patient had completed multiple courses of fluconazole without clinical improvement. Physical examination revealed no abnormalities. Rectal examination revealed a non-tender grade II prostate. Urinary tract ultrasound showed no significant abnormalities, with a structurally normal bladder and a postvoid residual urine volume of 20 ml. Prostate ultrasound revealed a homogeneous prostate with an approximate volume of 66 ml and overgrowth of the prostatic median lobe into the bladder (**Figure 1B**).

**Figure 1.**
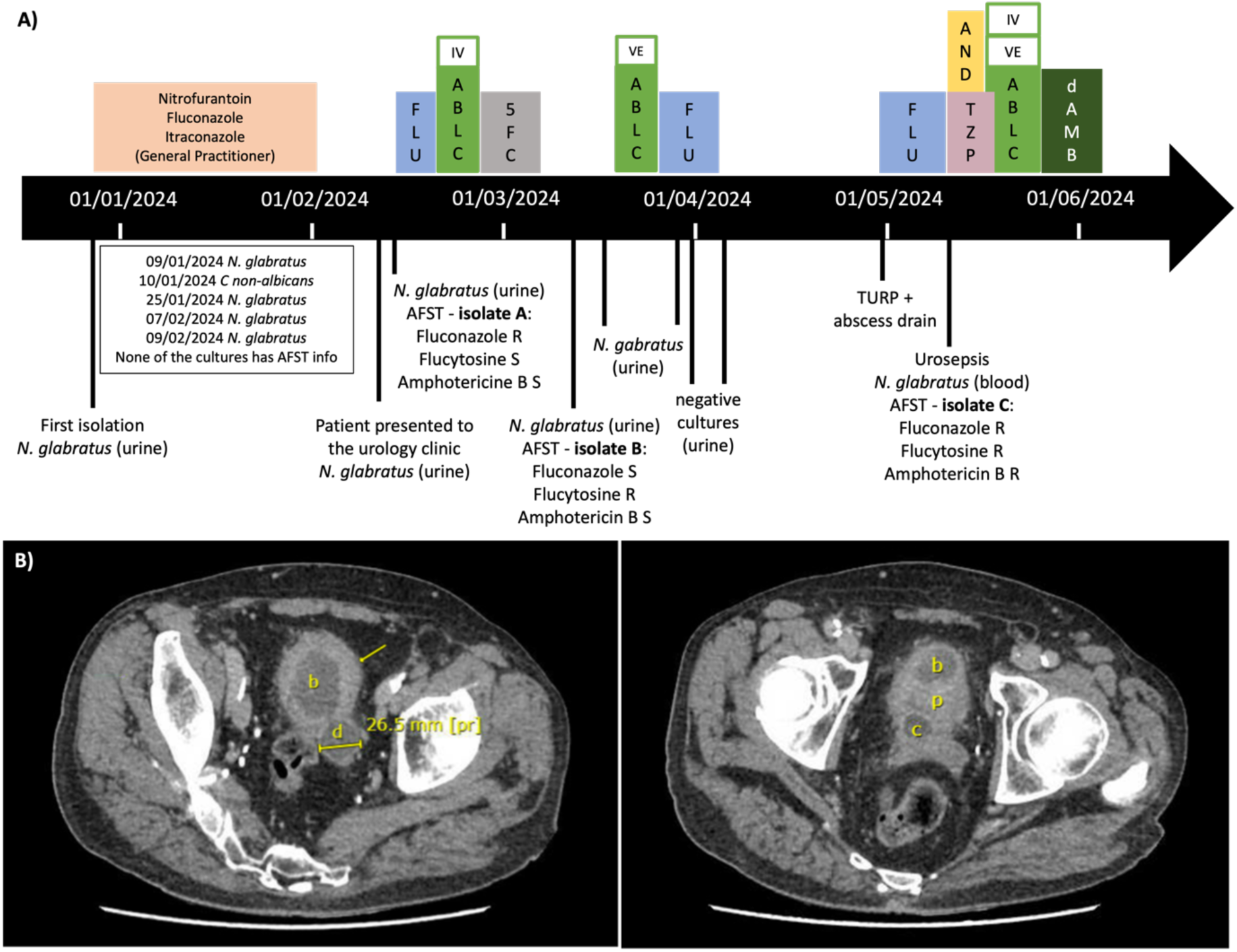
Timeline of isolation of *N. glabratus* and clinical manifestation. **A)** The timeline shows the isolation of strains and AFST data (below arrow) and treatment of the patient (above arrow). AFST details can be found in **Table 1**. FLU: fluconazole, ABLC: liposomal amphotericin B, VE: intravesical, IV: intravenously, 5FC: flucytosine; AND: anidulafungin, TZP: piperacillin-tazobactam; d-AMB: amphotericin B deoxycholate, TURP: transurethral resection of the prostate. AFST: antifungal susceptibility test. **B)** Abdominal and pelvic CT scan in axial section with enhancement. Left image: Bladder (b), bladder diverticulum (d) and thickened bladder wall (arrow). Right image: Bladder (b), prostate (p) and cystic structure (c).

**Table 1.** Clinical isolates with corresponding isolation date, site of infection, and AFST information if obtained. Interpretation was performed according to the clinical breakpoints as determined by CLSI for Sensititer YeastOne 10 (14) and EUCAST for the EUCAST reference method (15). N.A. indicates no available cl <u>inical breakpoint.</u>

| AFST |  |  |  |  |  |
| --- | --- | --- | --- | --- | --- |
| isolate | Date & AFST method | Site | Drug | MIC (µg/ml) | Interpretation |
| A | 14/02/2024<br>Sensititer YeastOne 10<br>(Thermo Fisher) | urine | Anidulafungin | 0.03 | Susceptible |
|  |  |  | Micafungin | 0.03 | Susceptible |
|  |  |  | Caspofungin | 0.5 | Resistant |
|  |  |  | 5-flucytosine | 0.06 | N.A. |
|  |  |  | Posaconazole | >8 | N.A. |
|  |  |  | Voriconazole | 4 | N.A. |
|  |  |  | Itraconazole | >16 | N.A. |
|  |  |  | Fluconazole | >128 | Resistant |
|  |  |  | Amphotericin B | 1 | Susceptible |
| B | 11/03/2024<br>Sensititer YeastOne 10<br>(Thermo Fisher) | urine | Anidulafungin | 0.015 | Susceptible |
|  |  |  | Micafungin | 0.015 | Susceptible |
|  |  |  | Caspofungin | 0.12 | Susceptible |
|  |  |  | 5-flucytosine | 64 | N.A. |
|  |  |  | Posaconazole | 0.25 | N.A. |
|  |  |  | Voriconazole | 0.25 | N.A. |
|  |  |  | Itraconazole | 0.5 | N.A. |
|  |  |  | Fluconazole | 4 | Susceptible<br>dose dependent |
|  |  |  | Amphotericin B | 1 | Susceptible |
| C | 10/05/2024<br>EUCAST | blood | Anidulafungin | <0.008 | Susceptible |
|  |  |  | Micafungin | <0.008 | Susceptible |
|  |  |  | 5-flucytosine | >32 | Resistant |
|  |  |  | Posaconazole | >8 | Resistant |
|  |  |  | Voriconazole | >8 | Resistant |
|  |  |  | Itraconazole | >8 | Resistant |
|  |  |  | Fluconazole | >128 | Resistant |
|  |  |  | Amphotericin B | 2 | Resistant |

Upon admission, the patient received fluconazole (400 mg daily, orally) for 5 days. The initial urine culture identified *N. glabratus* resistant to fluconazole but susceptible to flucytosine and amphotericin B (isolate A, **Table 1**). Consequently, fluconazole was discontinued, and amphotericin B lipid complex (670 mg daily, intravenously) was administered for 5 days. Treatment was subsequently switched to flucytosine (1500 mg four times per day, orally) for a duration of 10 days. Despite 1 week of flucytosine therapy, the patient continued to experience dysuria and pollakiuria. Cystoscopy revealed a bladder diverticulum with purulent drainage. A repeat urine culture identified *N. glabratus* resistant to flucytosine but susceptible to fluconazole and amphotericin B (isolate B, **Table 1**). Intravesical instillations of amphotericin B lipid complex (50 mg daily) were initiated, resulting in clinical improvement after 6 days. Fluconazole (400 mg daily, orally) was reintroduced for 8 days, leading to negative urine cultures. Unfortunately, symptoms recurred. A triple-phase abdomen and pelvis computed tomography (CT) scan revealed a thickened bladder wall, the previously known bladder diverticulum, and prostatic hypertrophy. Additionally, a 2 cm cystic structure was detected within the prostate (Figure 1B). No evidence of fungus balls was found. The patient underwent a transurethral prostatectomy, during which purulent drainage and calcified tissue debris were collected from the cyst. Treatment with fluconazole (800 mg daily, orally) was initiated. Histological analysis revealed fungal structures in fibrinpurulent material from TURP and acute and chronic inflammation of the prostatic urethra with no evidence of malignancy (**Figure S1,** Supplementary). 11 days after the surgery, the patient presented to the emergency department with acute anuria caused by clot retention. A complete blood count showed leukocytosis with a white blood cell count of 14 600/μL (93% neutrophils), a decrease in hemoglobin (Hb) to 8.4 g/dL (baseline ∼10 g/dL), and a normal to slightly elevated platelet count (432 000/μL). Serum creatinine increased significantly from the patient’s baseline of 1.3 mg/dL to 3.1 mg/dL, and C-reactive protein was markedly elevated at 160 mg/L. A single dose (1 250 mg) of amikacin was administered, and empirical therapy with piperacillin-tazobactam (4.5 g intravenously every 6 hours) and anidulafungin (200 mg loading dose, followed by 100 mg daily for 3 days, intravenously) was initiated. Fluconazole was discontinued. Patient showed no improvement after 48 hours. Blood and urine cultures were positive for *N. glabratus*, which was resistant to flucytosine, fluconazole and amphotericin B (isolate C, **Table 1**). Treatment with ABLC was initiated via both intravenous (3 mg, daily) and intravesical (50 mg, daily) routes for 11 days. Meanwhile, the pharmaceutical team submitted an import request for amphotericin B deoxycholate, which was expected to achieve higher urine concentrations but was not available in Belgium. Subsequently, amphotericin B deoxycholate treatment (25 mg daily, intravenously) was initiated and maintained for 10 days. Follow-up in the urology clinic revealed significant clinical improvement, and subsequent urine cultures remained sterile six months post-treatment.

### Genome analysis and sequence dataset mining

To investigate the mechanisms of acquired resistance to amphotericin B and flucytosine, we performed whole genome sequencing and variant calling analysis of the clinical strain using the ATCC2001 (CBS138) genome as a reference. To identify recently acquired variants that might explain the observed levels of resistance, we performed a phylogenomic analysis to determine the placement of this strain in the alignment of 420 *N. glabratus* isolates of Schikora-Tamarit & Gabaldón (16). Tree reconstruction revealed that the clinical isolate belongs to the *N. glabratus* clade 24 (16). A subset of the strains of this clade were previously also described as clade 7 (17). To differentiate clade-specific polymorphisms from potentially novel resistance-associated variants, we filtered out common variants present in 20% or more of the 51 strains within clade 24 (86713 variants) from the variant calling results of the clinical strain. This yielded a list of 73 protein-altering (missense, frameshift or nonsense) variants across 65 genes, listed in **Table S1** (Supplementary). Based on the functional annotation of the identified genes, we identified 4 mutations that can be linked to the resistance of this isolate, presented **Table 2**.

**Table 2.**
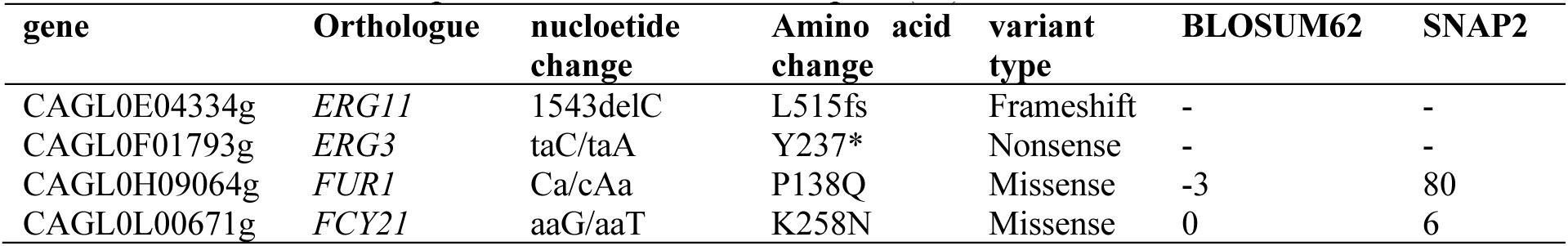
List of protein-altering variants in genes associated with antifungal drug resistance. found in the clinical isolate C. Given are gene IDs in *N. glabratus* reference genome (ATCC2001/CBS138), gene aliases or orthologous gene names in *S. cerevisiae*, nucleotide change indicating either specific variation in affected codons (reference allele/mutated allele) for single nucleotide polymorphisms (SNPs) or the nucleotide position in CDS for insertions (ins) or deletions (del). Predicted impacts of missense mutations on protein conformation were evaluated with **SNAP2** (18) and BLOSUM62 (19). A higher SNAP2 score suggests greater impacts on protein structure and function. BLOSUM62 scores <0 indicate a low likelihood of an amino acid substitutions to fixate by chance and can be used as a predictor for functional impact (20).

The *ERG3* and *ERG11* genes encode essential components of the ergosterol biosynthesis pathway. *ERG11* encodes lanosterol 14α-demethylase, which catalyses the demethylation of lanosterol, and *ERG3* encodes a C-5 sterol desaturase. The frameshift mutation in *ERG11* and nonsense mutation in *ERG3* (**Table 2**) suggest the LoF of these genes. LoF mutations in *ERG11* result in the accumulation of toxic sterols, which are normally converted downstream to ergosterol. However, simultaneous LoF mutations in *ERG3* block this conversion and prevent toxic sterol accumulation. The simultaneous LoF of *ERG3* and *ERG11* confers resistance to both amphotericin B and azoles, as ergosterol depletion reduces target binding, while also preventing toxic effects of sterol intermediates, preserving viability (10, 12).

*FUR1* and *FCY21* are linked to flucytosine antifungal activity and resistance. *FUR1* encodes uracil phosphoribosyltransferase, which catalyses the conversion of 5-fluorouracil (5-FU) into the toxic metabolite 5-fluorouridine monophosphate (5-FUMP), a crucial step in 5FC activity. 5-FUMP inhibits both RNA and DNA synthesis, leading to cell death. Mutations in *FUR1* can prevent this conversion, rendering flucytosine ineffective (21). *FCY21*, a cytosine permease, facilitates the uptake of flucytosine into fungal cells. Mutations that impair *FCY21* activity can decrease flucytosine import, contributing to low-level resistance in *S. cerevisiae* (22). The high SNAP2 score and low BLOSUM62 score of the missense mutation in *FUR1* (**Table 2**) suggest a high functional impact and a rarely found amino acid substitution respectively. The opposite is true for *FCY21* (Table 2), suggesting this mutation might be of low impact and is likely to be the product of neutral genetic drift.

Because *FUR1* mutations have been investigated and associated with acquired flucytosine resistance to a great extent (21, 23), but the simultaneous LoF of *ERG3* and *ERG11* has not been reported in clinical isolates of *N. glabratus,* we further focussed on the latter mechanism of resistance for further investigation.

We investigated whether variation in both *ERG3* and *ERG11* is common in *Candida* strains, by screening a dataset of genomes of *C. albicans* (642 strains), *C. auris* (752 strains), *N. glabratus* (420 strains), *Candida orthopsilosis* (33 strains), *C. parapsilosis* (51 strains), and *C. tropicalis* (89 strains) reported by Schikora-Tamarit & Gabaldón (16). Only protein-altering variants such as stop-gained, frameshift, splice-site in-frame insertions/deletions, and missense variants with a frequency lower than 0.2 (<20%) were considered in this analysis. For *N. glabratus* and *C. parapsilosis* we did not identify cases of co-occurrence of *ERG3* and *ERG11* variants in the filtered clinical dataset. In contrast, we identified isolates with co-altered genes for *C. auris* (1), *C. orthopsilosis* (1), *C. tropicalis* (6) and *C. albicans* (73), as shown in **Table S2** (Supplementary). The relatively high frequency of concerted *ERG3/ERG11* mutations in *C. albicans* was surprising. Upon closer inspection, we found out that the majority of strains carried often similar combinations of mutations among which A351V and/or A353T mutation in *ERG3* gene, along with *ERG11* variants such as E266D, V437I, S442F and V488I, and similar combinations were found enriched in the same clades. Among all analyzed *C. albicans* strains, only one strain demonstrated a rare combination of mutations in both genes: *ERG3* D14N, identified in one strain, and *ERG11* E336G, present in 3.27% of analysed strains (21 strains). This suggests that most of such combinations might not represent recently-emerged clinical adaptations but rather pre-existing variation present within specific clades that passed our 20% threshold at the species level. Similarly, of the six concerted *ERG3/ERG11* mutations, five were present in environmental strains underscoring they are naturally occurring non-synonymous variations, unrelated to clinical adaptation.

Additionally, all identified co-occurring mutations were missense variants, while the clinical isolate of this study shows a frame-shift and nonsense mutation in *ERG11* and *ERG3* respectively (**Table 2**). Overall, the combination of high SNAP2 scores (>50) and low BLOSUM62 values (<1) for both *ERG11* and *ERG3* missense mutations, were not detected. This suggests that high-impact variation in both *ERG3* and *ERG11* genes is rare among *Candida* species. Consistently, we found no reports in the literature of combined LoF mutations in both *ERG3* and *ERG11* in clinical strains of *N. glabratus*.

### Validation of the mechanism of amphotericin B resistance

We further investigated the effects of the LoF of *ERG3* and *ERG11*, by constructing single and double deletion mutants of both genes in the ATCC2001 (CBS138) background, followed by antifungal susceptibility testing and sterol analysis. The deletion of *ERG11* in the WT background could not be achieved in transformations under standard conditions, but was successful after plating transformed cells on agar supplemented with ergosterol and Tween 80 and incubating them anaerobically at 37 °C, following recommendations from Geber *et al.* (11). The deletion of *ERG11* in the *ERG3Δ* background was achieved in standard transformation conditions. Attempts to delete *ERG3* and *ERG11* in the clinical strain background were unsuccessful, potentially due to the high stress sensitivity (see further), poor growth and short life span of the isolate. During experimentation, we noted that when grown in RPMI-MOPS 2% glucose liquid medium at 37°C, a significant amount of *ERG1111, ERG311+ ERG1111* and clinical strain (C1 and C2) cells die, as shown by LIVE/DEAD staining at 0h and 24h post-incubation (see **Figure S2**, Supplementary).

Broth dilution assays (BDA) showed that the LoF of *ERG11* lead to amphotericin B and azole (fluconazole, posaconazole and ketoconazole) resistance and resulted in collateral sensitivity to echinocandins (caspofungin, anidulafungin and micafungin) (Figure 2A). Simultaneous LoF of *ERG3* and *ERG11* yielded susceptibility patterns similar to the *ERG11* LoF strains. Interestingly, the *ERG311+ERG1111* strains showed a 4-fold increase in flucytosine resistance, based on ETEST results (Figure 2B). Nevertheless, this reduced susceptibility cannot explain the hyper-resistance for flucytosine in the clinical strain, which is likely the result of additional mutations in *FUR1, FCY21* and/or other genes. *ERG3Δ* did not show altered susceptibility to amphotericin B, echinocandins or flucytosine but did show slight changes in azole susceptibility. The MIC decreases for all azoles, while the higher supra-MIC growth was detected for ketoconazole.

**Figure 2.**
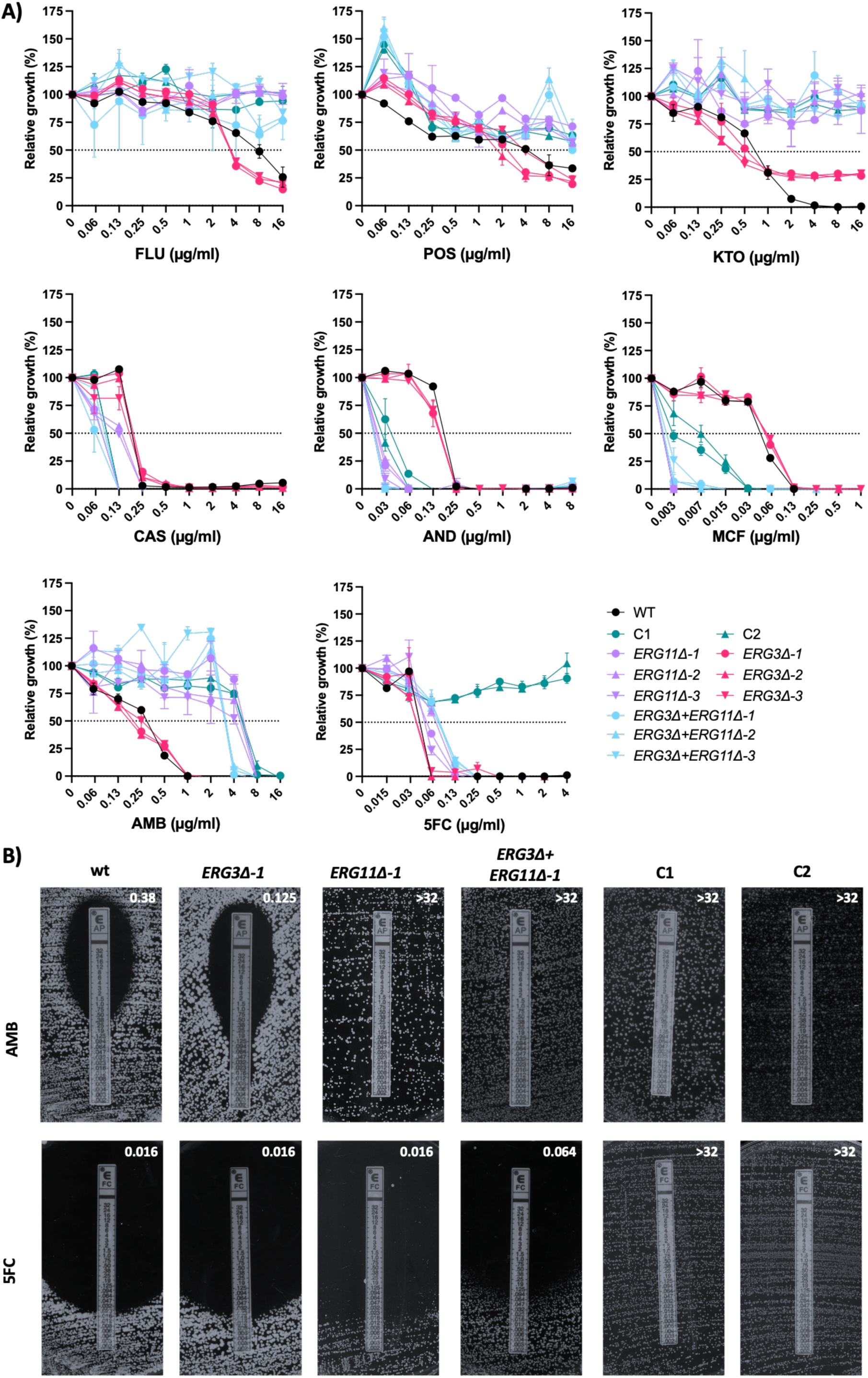
Antifungal susceptibility testing (AFST) of clinical and constructed strains. **A)** BDA for fluconazole (FLU), posaconazole (POS), ketoconazole (KTO), caspofungin (CAS), anidulafungin (AND), micafungin (MCF), amphotericin B (AMB) and flucytosine (5FC) for the WT *N. glabratus* (strain ATCC2001) and 3 biological replicates (independent trasformants) per constructed genotype (i.e. *ERG3Δ, ERG1111* and *ERG311+ ERG11Δ*) in the WT background. Error bars indicate standard deviation based on two technical replicates. C1 and C2 are two single-colony isolates from the clinical isolate C. **B)** pictures of ETEST® (bioMérieux) after 72 hours incubation at 37 °C. The MIC value (in µg/ml) is read as the lowest concentration at which the border of the elliptical growth inhibition zone intercepted the strip, and is indicated in the top-right corner of each image.

Next, we performed relative total sterol quantification on stationary growth phase cells from all constructed and clinical strains, following Carolus *et al.* (9). The analysis shown in Figure 3 indicates that ergosterol, which is dominant in the WT strain, is replaced mainly by 4,14-dimethyl-zymosterol and lanosterol in the *ERG311+ ERG1111 -* strains and in the clinical strains (C1 and C2), confirming that Erg3 and Erg11 are non-functional in the clinical isolate. As expected, the *ERG1111* strains show the accumulation of toxic 14-methyl-ergosta-8,24(28)-diene-3,6-diol, which is absent from *ERG311+ERG1111* -or clinical strains. The deletion of *ERG3* leads to domination of ergosta-7,22-dienol and episterol, which still might bind amphotericin B, as suggested by data presented in Figure 2.

**Figure 3.**
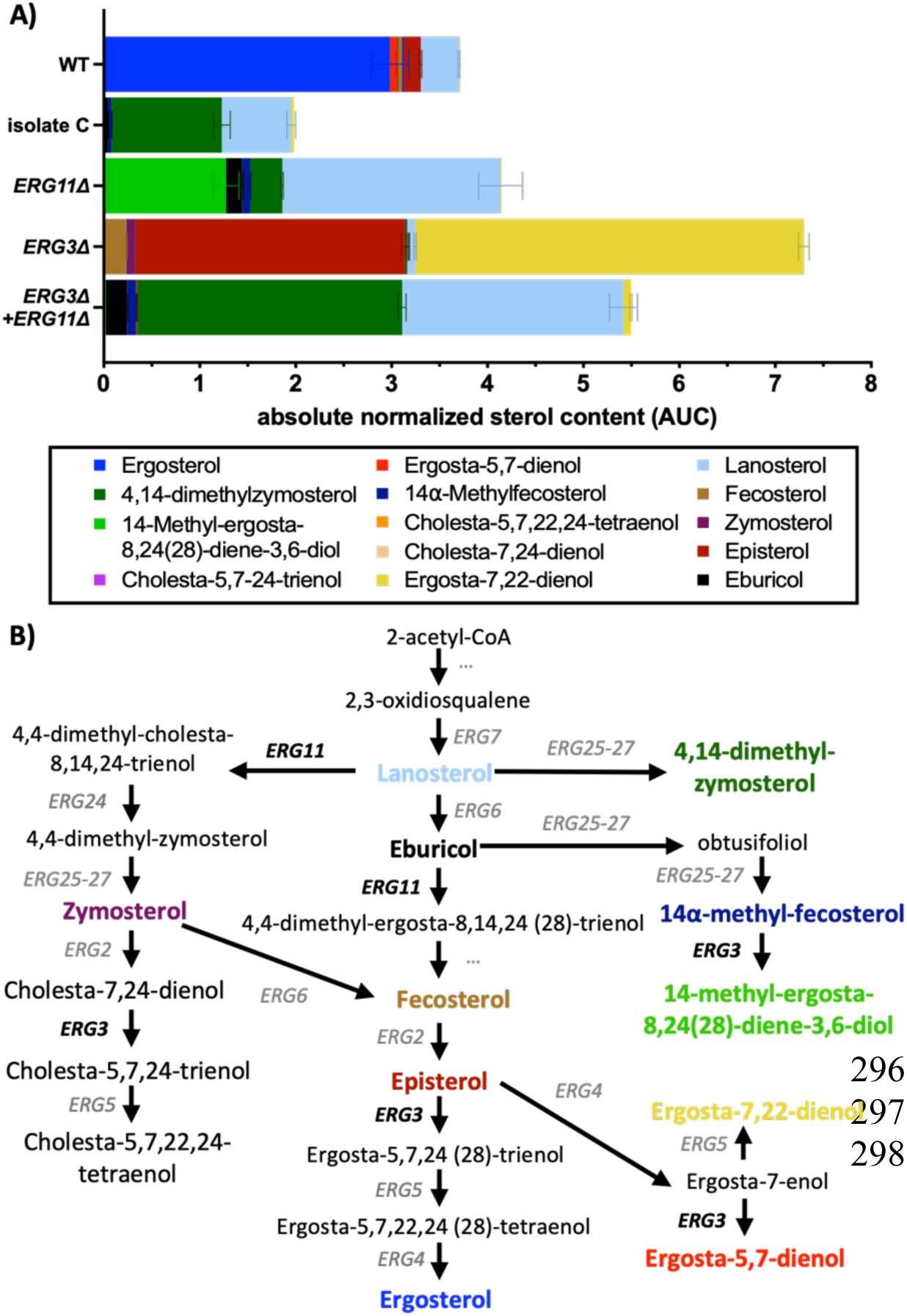
Sterol quantification. **A)** Bar plots show sterol abundance relative to cholestane as an internal standard, and are based on the area under the curve (AUC) of the GC-MS response. Cells are grown in drug-free growth conditions. Error bars indicate standard deviation from a minimum of two technical repeats (independent culture and GC–MS run) for one representative strain. **B)** Simplified representation of the ergosterol biosynthesis pathway. Components and genes in bold are the main affected sterols and genes in our sequenced strains. The colours match the sterol compounds in the bar plots of A).

### Investigation of fitness trade-offs

Due to the notable slow growth and short life-span of the clinical isolates, and several constructed strains, we performed a growth and stress susceptibility analysis of all strains to estimate the burden of the LoF of *ERG3* and/or *ERG11* on cellular fitness. Growth curves shown in Figure 4A demonstrate a strong reduction in growth rate, carrying capacity and increase in lag-phase, due to the LoF of *ERG11* or *ERG3* and *ERG11,* in most growth conditions. The clinical strains seem to be slightly more adapted to low-temperature conditions (30°C) in physiological media (RPMI-MOPS 0.2% glucose and 2% glucose), compared to the *ERG311+ERG1111* and *ERG1111* strains. The LoF of *ERG3* does not significantly affect growth in most conditions, except in RPMI-MOPS 2% glucose, in which they grow significantly less than the WT strain at 30°C but better than the WT strain at 41°C. A similar trend is observed in other media, but to a lesser extent. Thus, an *ERG3Δ* seems to impact thermotolerance in *N. glabratus.* The LoF of *ERG11* on the other hand, renders any strain inviable at high febrile temperatures, as no growth was recorded for any *ERG11Δ, ERG311+ERG11Δ* or clinical strain at 41°C in any of the growth media.

**Figure 4:**
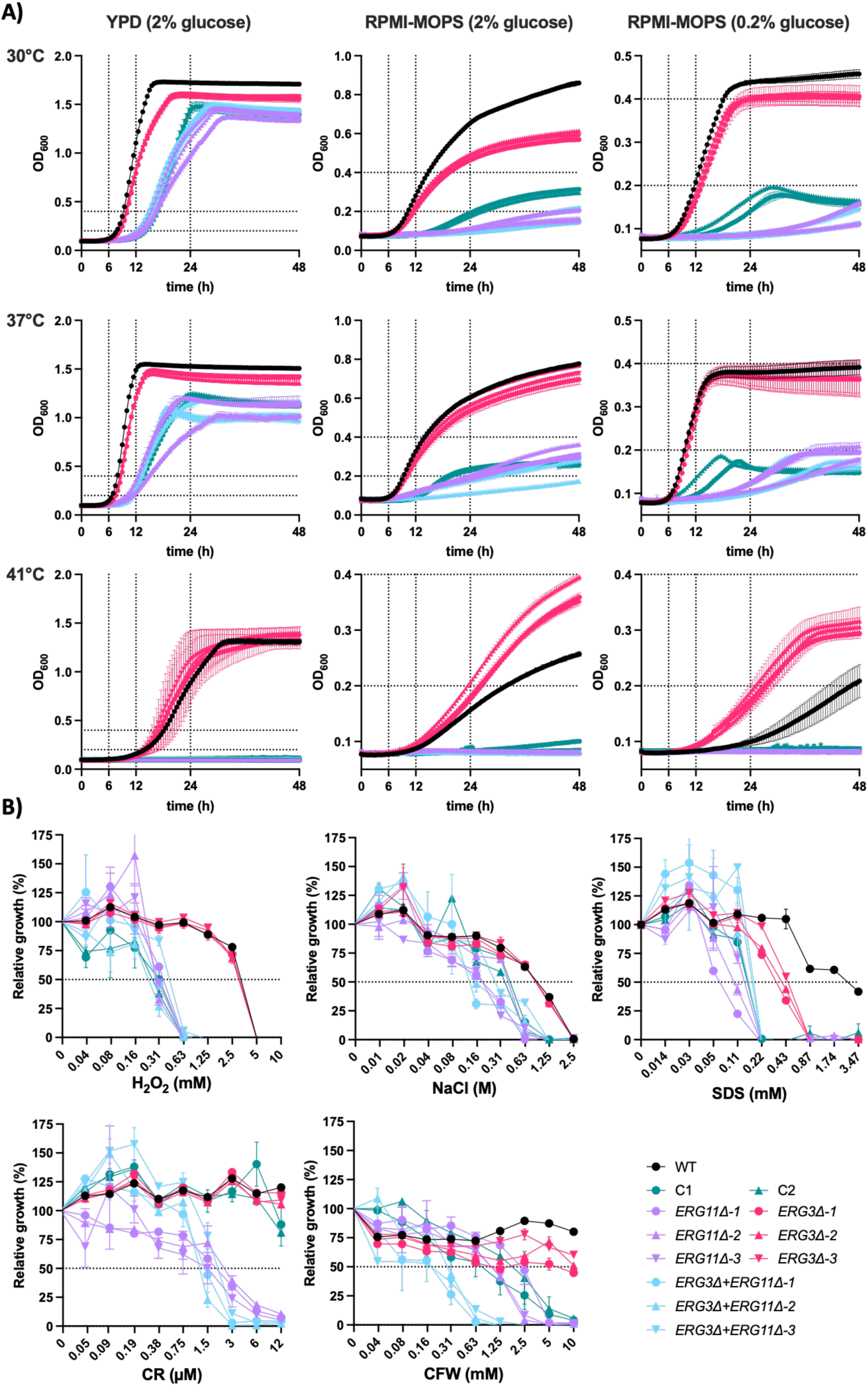
Fitness profiling of clinical and constructed strains. **A)** growth evaluation over 48h at three different temperatures (30°C, 37°C and 41°C) and in three different media (YPD+2% glucose, RPMI-MOPS+0.2% glucose and RPMI-MOPS+2% glucose). Colour and symbol legends for different strains are given in B. Error bars indicate standard deviation based on two technical repeats for each strain. **B)** Broth dilution assays (BDA) depict the relative growth as a function of stressor concentration in RPMI-MOPS (pH 7, 2% glucose) after 48h of incubation. The stress-inducing compounds used were calcofluor white (CFW), Congo red, (CR), sodium dodecyl sulfate (SDS), sodium chloride (NaCl) and hydrogen peroxide (H_2_O_2_). Error bars represent the standard deviation (SD) of two technical repeats per strain.

Similar to growth trends, stress susceptibility assays show that the LoF of *ERG11* but not *ERG3* significantly impacts fitness, as shown by the tolerance profiles for most stressors in Figure 4B. Membrane stress tolerance (exerted by the detergent sodium dodecyl sulfate, SDS) is reduced to a lesser extent in *ERG3Δ* compared to *ERG11Δ* or *ERG311+ERG1111* strains, indicating that membranes enriched in episterol and ergosta-7,22-dienol are less stable than ergosterol enriched membranes, but more stable or functional than those enriched with lanosterol, 4,14-dimethyl zymosterol and/or 14-methyl-ergosta-8,24(28)-diene-3,6-diol (Figure 3). The tolerance to oxidative stress (hydrogen peroxide susceptibility) and osmotic stress (sodium chloride susceptibility), is reduced similarly for the *ERG1111, ERG311+ERG11Δ,* and clinical strains. Remarkably, the collateral sensitivity to the cell wall stressors congo red and calcofluor white, seen in the *ERG1111* and *ERG311+ERG1111* strains, is not present or present to a lesser extent in the clinical strains, suggesting that additional mechanisms or the strain background plays an important role in cell wall stress resistance. The absence of glucan-binding stress exerted by congo red in the clinical strain, might explain the lower susceptibility to micafungin seen in Figure 2A.

In summary, the LoF of *ERG11* severely impacts fitness in terms of growth capacity and stress tolerance, while some of this reduced fitness is mitigated by the LoF of *ERG3* and/or background specific differences.

### Nitroxoline as therapeutic opportunity with novel mechanism of action?

With poor urinary penetration of echinocandins (4, 5), the high resistance to amphotericin B, flucytosine and azoles in this UTI case, leaves few therapeutic options. Recently, the antibiotic nitroxoline (8-hydroxy-5-nitroquinoline) which shows high urinary excretion, was reported to have antifungal activity against several *Candida* species, including *N. glabratus* (24, 25) and multidrug resistant *C. auris* (26, 27). Therefore, we tested the susceptibility towards this compound, and found that the clinical strain has a low MIC (0.5 µg/mL), while *ERG311, ERG1111* and *ERG311+ERG11Δ* exhibit collateral sensitivity, compared to the WT strain (Figure 5A). The iron chelating properties of the 8-hydroxyquinoline group, which is a privileged structure (28), is hypothesized to exert its antimicrobial activity, although other modes of action have been proposed (29, 30). To further evaluate whether, the chelating properties indeed exert the antifungal and collateral sensitive effect of nitroxoline on our tested strains, we evaluated the susceptibility towards the drugs ciclopirox and diiodohydroxyquinoline, and towards the iron chelator bathophenanthroline disulfonic acid (BPS). Ciclopirox is a topical antifungal of the hydroxypyridone class, which exerts its activity through iron chelation (31). Diiodohydroxyquinoline (8-Hydroxy-5,7-diiodoquinoline or Iodoquinol) is an antiprotozoal drug, which also exerts antifungal activity and it is, like nitroxoline, an 8-hydroxyquinoline derivative (32). Figure 5A shows that the response to other iron chelating agents (BPS and ciclopirox) or the 8-hydroxyquinoline analogue diiodohydroxyquinoline is significantly different from the one towards nitroxoline. The LoF of neither *ERG11*, nor *ERG3,* seems to affect the tolerance towards iron chelation, while the clinical strains show a higher MIC towards diiodohydroxyquinoline, compared to the WT and constructed strains. This suggests that the antifungal activity of nitroxoline extends beyond the iron chelating properties of its 8-hydroxyquinoline group.

**Figure 5.**
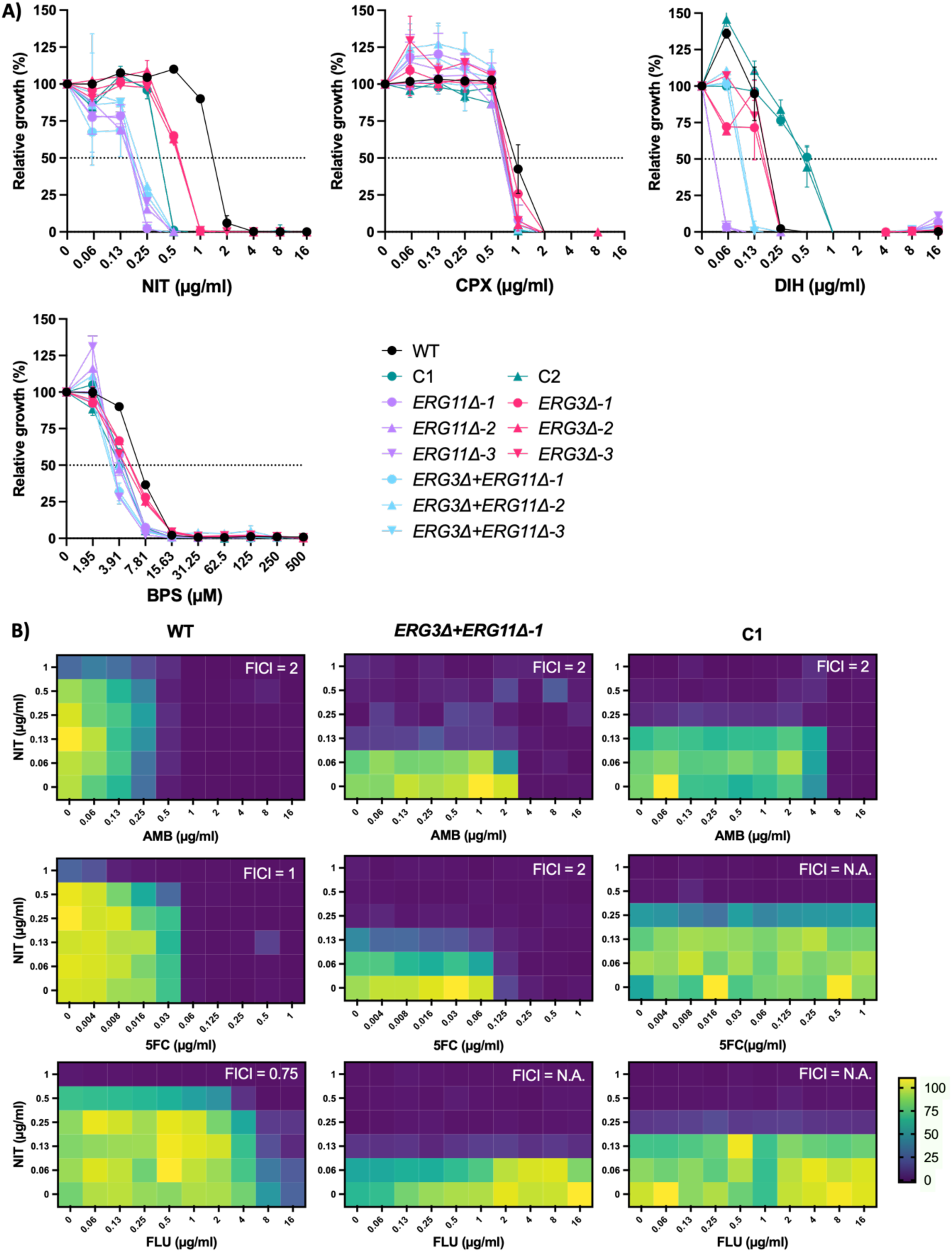
Evaluation of nitroxoline and compounds. **A)** Broth dilution assays (BDA) depicted as the relative growth in function of compound concentration in RPMI-MOPS (pH 7, 2% glucose) after 48h of incubation. Compounds used are nitroxoline (NIT), ciclopirox ethanolamine (CPO), diiodohydroxyquinoline (DIH), and bathophenanthrolinedisulfonate (BPS). Error bars represent the standard deviation (SD) of two technical repeats per strain. **B)** Checkerboard assays to evaluate drug interactions between NIT and amphotericin B (AMB), flucytosine (5FC) and fluconazole (FLU) respectively for the WT, *ERG311+ERG1111* and clinical (C1) strain. To evaluate drug interactive effects, the fractional inhibitory concentration index (FICI) was calculated and indicated in the right upper corner. 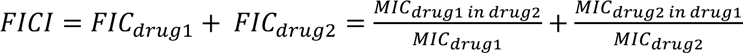, with *MIC_drug 1 in drug 2_* being the MIC of drug 1 in the presence of the highest concentration of drug 2 where growth is still visible (>50% compared to control without drug). FICI ≤ 0.5 indicates synergy, 0.5< FICI <1 indicates an additive effect, 1≤ FICI < 4 indicates indifference and a FICI > 4 indicates antagonism (33). For checkerboard assays in which the MIC of one of the two drugs could not be determined within the tested range, no FICI could be calculated (indicated by N.A.).

Finally, we evaluated the interactive effects of nitroxoline with conventional antifungals— fluconazole, amphotericin B, and flucytosine—with checkerboard assays, to determine whether this repurposed drug exhibits synergistic or antagonistic antifungal activity. Figure 5B shows that nitroxoline does not increase the activity of any of the drugs towards the WT, *ERG311+ERG1111* or clinical strain. The fluconazole – nitroxoline combination showed an additive effect with a fractional inhibitory concentration index (FICI) of 0.75, while all other drug combinations were indifferent or 1≤ FICI < 4 (33), or could not be calculated because the high resistance to the drug under investigation did not yield an MIC result within the tested range. Thus, while nitroxoline shows great potential as a repurposed antifungal agent for the treatment of complicated MDR UTI’s like in this case, the combination of nitroxoline with fluconazole, amphotericin B or flucytosine, does not seem to increase its therapeutic potential.

## Discussion

This study uncovers a rare case of multidrug resistant *N. glabratus* with dysfunctional Erg3 and Erg11 and severely impaired fitness that emerged under prolonged antifungal therapy in a complicated and persistent UTI that led to a breakthrough candidemia. LoF mutations in *ERG3* and *ERG11* resulted in resistance to amphotericin B, azoles and decreased sensitivity to flucytosine, while mutations in *FUR1* and/or *FCY21* likely conferred high flucytosine resistance in the clinical isolate. Since polyenes, azoles, and flucytosine encompass most of the antifungal arsenal for treating UTIs, we explored the urinary tract antibiotic nitroxoline, which our results show to be a potent therapeutic alternative that may exert its antifungal activity via an unknown mechanism, unrelated to iron chelation.

Prior work has shown that *ERG3* or *ERG11* defects individually can drive azole and/or amphotericin B resistance in *Candida* species (10, 34, 35), but the combined LoF of both genes is extremely rare, as shown by our *Candida* genome dataset screening. After one *C. tropicalis* case from a Tunisian hospital reported by Eddouzi *et al*. (13), this is the second case in which both *ERG3* and *ERG11* LoF mutations confer azole and amphotericin B cross resistance in a *Candida* strain from clinical origin. It is the first report of this mechanism of resistance in *N. glabratus.* In addition, the co-occurrence of hyper-resistance to flucytosine within this background is unique, and we hypothesize it is the result of additional mutations in *FUR1* and/or *FCY21,* which have both been implicated in flucytosine resistance (21–23).

The *ERG311+ERG11Δ* genotype has been investigated in several experimental evolution and gene knock-out studies (7–9, 11, 12). Our sterol analysis demonstrated that the LoF of *ERG3* and *ERG11* in *N. glabratus,* lead to a 4,14-dimethyl-zymosterol and lanosterol dominated sterol profile, which is similar to the effect observed in *C. albicans* (12) and *C. auris* (9), but different from the profile observed in *C. tropicalis,* which was dominated by 14α-methyl fecosterol, obtusifoliol and eburicol (13). Another study in *N. glabratus* also shows 14α-methyl fecosterol, and obtusifoliol, alongside lanosterol and 4,14-dimethyl-zymosterol (11). A potential explanation might be that the sterol analyses in this study and the study of *C. albicans* (12) and *C. auris* (9), were performed on stationary phase cells or saturated cultures, while the analyses on *C. tropicalis* (13) and *N. glabratus* (11) were performed on cells in exponential growth phase (overnight culture) (13), which could result in the detection of short-lived intermediate sterols. Alternatively, significant differences could exist in sterol biosynthesis pathways between strains and species. The latter is supported by the observation that the *ERG311* genotype leads to the accumulation of ergosta-5,7-dienol in this study, while this intermediate is absent in the same genotype in *C. auris,* grown and analysed under the same conditions (9). Similarly, the LoF of *ERG3* has been reported to display resistance to azoles and cross-resistance to polyenes in *C. albicans* (8, 35) and *C. auris* (9, 36), while other studies and our results show no effect or even an increased susceptibility to either or both drug classes in the *ERG311* background (11, 12). This, along with the fact that high impact *ERG3* variation is rare (34), which is confirmed by our clinical dataset screening, suggests that *ERG3* variation is no major driver of resistance. In our case, the LoF of *ERG3* is a mechanism to compensate for the deleterious LoF of *ERG11,* rather than a stand-alone mechanism of polyene and/or azole resistance. The LoF of *ERG3* partially rescues several fitness trade-offs caused by the LoF of *ERG11,* as shown by our results. Fitness trade-off compensation in antifungal drug resistance evolution has recently been first observed in acquired amphotericin B resistance in *C. auris,* and such mechanisms are hypothesized to drive resistance *in vivo* (9).

Because most types of sterol modulation that confer amphotericin B resistance impose significant fitness costs, it has been proposed that these changes reduce virulence and restrict the evolution of such mechanisms *in vivo*, making them less likely to occur in infections (8, 9). The simultaneous LoF of *ERG3* and *ERG11* has been reported to reduce the growth rate and stress tolerance significantly (8, 9, 11), to the extent that cells were assumed to be no longer able to maintain a bloodstream infection in an *in silico* infection model for *C. auris* (9). The fact that this genotype was isolated from a blood infection, and it showed hyper-resistance to flucytosine by an additional mechanism, challenges the notion that fitness trade-offs always undermine the evolution of resistance *in vivo.* This study shows that the imbalance of stronger drug resistance coupled with impaired growth, short life-span and high stress sensitivity as a cost, does not hinder in-host adaptation. Although the selective pressure of antifungal treatment and/or the infection site (the prostate is a pharmacologic sanctuary site) may favour severely compromised mutants, the breakthrough candidemia of this isolate shows that these strains can persist and cause recalcitrant infections of other niches. It is important to note though, that besides ‘isolate C’, no other clinical isolates were stored or could be investigated. Whether the *ERG3* and *ERG11* LoF mutations were acquired in the UTI or later, during candidemia, is therefore unknown.

Due to the limited arsenal of antifungal drugs that can be used in fungal infections of the urinary tract in which resistance to polyenes, azoles and flucytosine is acquired, we investigated the *in vitro* potency of nitroxoline, an antibiotic with high urinary excretion and antifungal activity (24–27). All strains were susceptible to nitroxoline and additionally, the *ERG3Δ, ERG1111* and *ERG311+ERG1111* strains showed collateral sensitivity to this drug. Collateral sensitivity is the phenomenon by which a strain acquires increased sensitivity to one drug due to the acquisition of resistance to another drug (here collateral sensitivity to nitroxoline due to resistance to azoles/polyenes). This phenomenon is well studied in bacteriology and oncology but has only recently been systematically studied in fungi, where collateral sensitivity-based drug cycling was shown to both prevent and actively reduce antifungal drug resistance (27). Besides drug cycling, combination therapy in which collateral sensitive drug pairs are used, can be an effective therapeutic improvement. Drugs that cause collateral sensitivity and interact in synergy – when the combined effect of two drugs is greater than their individual effect – would be good candidates for combination therapy (37–40). We did not identify a synergistic interaction between nitroxoline and fluconazole, amphotericin B or flucytosine, although we also did not detect antagonism. Nevertheless, even if drug pairs are solely additive or even indifferent or antagonistic, they can reduce resistance development, as the type of interactions between antimicrobials — whether synergistic, additive, or antagonistic — does not correlate to evolutionary outcome during combination treatments in bacteria, likely because these interactions can also evolve over time and collateral sensitivity can play a bigger role (39–41). Thus, the absence of synergy does not limit the potential of nitroxoline, as sole treatment or in combination with antifungals, in complicated UTI’s and we highly recommend further investigations for this purpose.

Nitroxoline is thought to exerts its antifungal activity primarily through its ability to chelate metal ions—particularly iron—thereby disrupting metal-dependent processes within the fungal cell. Vincent *et al.* detected collateral sensitivity towards the iron chelator BPS in *ERG311+ERG1111* strains of *C. albicans* (8). Surprisingly, we did not detect collateral or increased sensitivity towards BPS and ciclopirox, known iron chelating antifungal agents, in the constructed mutants and clinical strain. Additionally, diiodohydroxyquinoline, an antiparasitic 8-hydroxyquinoline derivative like nitroxoline, showed a different effect compared to nitroxoline, with lower susceptibility towards the clinical isolate, compared to the WT strain. These observations support the hypothesis that nitroxoline has additional modes of action, beyond iron chelation. Recently, nitroxoline was found to affect membrane integrity in bacteria (30) while in fungi, interactions with alkane 1-monooxygenase and methionine aminopeptidase enzymes were proposed, while no direct effects on cell membrane or cell wall were evidenced (29). Additional research into the mode of action and clinical applicability of nitroxoline as an antifungal compound are therefore highly desirable.

In summary, this case highlights that even profoundly “unfit” fungal strains can thrive under intensive antifungal therapy when hyper-resistance to multiple drugs is acquired, challenging the traditional premise that severe fitness costs necessarily preclude *in vivo* persistence under treatment. The promising *in vitro* efficacy and collateral sensitivity observed for nitroxoline against these isolates underscores its potential as a viable alternative agent when conventional treatments fail. Taken together, these findings emphasize the dynamic interplay between fitness trade-offs and resistance evolution and point to repurposed therapeutics—like nitroxoline—as essential antifungal agents for recalcitrant MDR fungal infections.

## Methods

### Ethical statement

All research presented in this study was conducted according to the ethical guidelines of the ethical committee of ZAS, UZ Leuven and KU Leuven. Informed consent was obtained from the patient and ethical approval was obtained from the ZAS committee for medical ethics (recognition number 009).

### Strains and growth conditions

Strains were stored at −80°C in 20% glycerol and routinely plated on solid YPD (1% yeast extract, 2% bacteriological peptone, 2% dextrose) agar (2%) at 37°C. Unless specified otherwise, cells were grown in MOPS (morpholinopropane sulfonic acid) buffered (pH 7) RPMI 1640 (Thermo Fisher Scientific) medium with 2% total glucose, at 37°C.

### Whole genome sequencing (WGS)

The clinical *N. glabratus* isolate was isolated from the patient and cultured on Sabouraud dextrose agar and incubated at 37°C for 48 hours. Genomic DNA was extracted using the ZymoBIOMICS DNA/RNA Miniprep Kit (Zymoresearch, USA), according to the manufacturer’s guidelines. The purified genomic DNA was quantified using a Qubit fluorometer. Library and sample preparation were carried out using the Nextera XT Sample Preparation Kit (Illumina, USA). Sequencing was performed on a MiSeq platform (Illumina Inc., USA) using 2 x 250 bp paired-end sequencing v2 500 cycles.

### WGS data analysis

Whole genome sequencing analysis was performed using the perSVade software (version 1.02.6) (42). To remove adaptors and trim the reads for each sample, we used FastQC (v. 0.11.9) (43) and Trimmomatic (v. 0.38)(44) with default parameters, facilitated by the trim_reads_and_QC module of perSVade. Alignment of the trimmed reads (with BWA MEM (v. 0.7.17) to the *N. glabratus* ATCC2001 (CBS138) reference genome (version s02-m07-r35), available on the Candida Genome Database (CGD), was performed with align_reads module of perSVade. Genome-wide coverage distributions were visualized using WGSCoveragePlotter, a component of the Jvarkit suite (v. c789c6a41 10.6084/m9.figshare.1425030, see **Figure S3**, Supplementary). After that, module call_small_variants integrates results of variant calling on the aligned reads (SNPs and small indels) from three different variant callers: BCFtools (v. 1.9)(45), GATK Haplotype Caller (v. 4.1.2)(46), and Freebayes (v. 1.3.1)(47).

For variant calling, we used the parameter ‘–ploidy 1’, under the assumption that the strain has the canonical ploidy. Variants with coverage below 12 and a minimum fraction of reads covering a variant (--min_AF) below 0.9, were filtered out. Moreover, obtained variants were filtered to retain only those that passed the filters of minimum two callers. Frameshift variants with coverage of more than 20 that did not initially meet the original mean allele frequency (min_AF) threshold but were supported by all three callers (lowering the min_AF threshold to 0.85), were included in the final set of variants. This approach mitigates tool-specific biases in indel calling, such as differences in local realignment and breakpoint variability across the used variant callers.

Next, we performed a phylogenetic tree reconstruction for the clinical strain of this study with 420 *N. glabratus* strains (393 of them have a clinical origin) studied in Schikora-Tamarit & Gabaldón (16). Using variant calling data for each strain we constructed a multi-sample VCF file and generated a pseudo-alignment using vcf2phylip tool. To avoid the biases introduced by indels, only homozygous SNPs were included in the pseudo-alignment. We then obtained the unrooted tree, using IQ-TREE (v2.1.2), from this pseudo-alignment using ‘-m TEST’ to use default automatic model selection. Ascertainment bias correction (+ASC) was not included in the model, as the primary objective of the phylogenetic tree reconstruction was not to estimate branch lengths, but to place the strain of interest within the pre-established phylogeny of strains and identify the most likely clade this strain belongs to. Next, we used midpoint rooting to obtain the final tree, which has support values from 1,000 bootstraps.

After determining the clade, all the members of this clade were used to create an artificial background, consisting of SNPs present in more than 20% of the clade members, to filter the variant calling results of the clinical strains and remove SNPs that were identified because of the difference of this strain with the reference genome used. Variants included in the artificial background were filtered out from the high-confidence sets of the clinical strain (with BCFtools v. 1.15.1(20) function isec). After that the annotation of final sets of variants was performed with annotate_small_vars module of Ensembl Variant Effect Predictor v. 100.2(24), incorporated in perSVade.

The filtering criteria for variant calling of the studied strain were selected to align with the established variant calling procedure used in Schikora-Tamarit & Gabaldón (16). This consistency ensures that the clinical strain was consistently and accurately placed within the reconstructed phylogenetic tree.

### Clinical variant data analysis

The co-occurence of protein-altering variation in both *ERG3* and *ERG11* was analyzed by mining the dataset of Schikora-Tamarit & Gabaldón (16). This dataset comprises variant calling results for 2000 genomes of *Candida albicans*, *Candida auris*, *Nakaseomyces glabratus* (*C. glabrata*), *Candida orthopsilosis*, *Candida parapsilosis*, and *Candida tropicalis*.

Protein-altering variants were identified based on Variant Effect Predictor (VEP) annotations generated in Schikora-Tamarit & Gabaldón (16).

The frequency of each variant was calculated as the proportion of strains with a variant relative to the total number of strains analyzed for each species. Variants in genes of interest were extracted using corresponding gene IDs for *ERG3* and *ERG11* for each species as annotated in the reference genomes of each species.

Only protein-altering variants, including stop-gained, frameshift, in-frame insertions/deletions, and missense variants with a frequency of less than 0.2 (<20%) were considered in this analysis. This approach aimed to eliminate background genetic variation characteristic of the species or clade, highlighting mutations that could represent recent or species-specific adaptations with potential phenotypic relevance.

### Strain construction

Deletion cassettes for *ERG3* (CAGL0F01793g) and *ERG11* (CAGL0E04334g), consisting of a nourseothricin resistance marker flanked by Flp recombinase recognition sites and 500 bp upstream and downstream homology regions of the target genes, were constructed in the pYC44 vector. The 500 bp up and downstream regions of *ERG3* and *ERG11* were amplified from the genomic DNA of the wild-type strain (ATCC2001) with primers designed to include an EciI recognition site at the cassette’s outer ends. Primers are specified in **Table S3** (Supplementary). The pYC44 vector was digested with XhoI and BamHI, and the promoter and terminator regions were inserted via Gibson Assembly. Correct insertion was verified by PCR using primers listed in **Table S3** (Supplementary). To generate linear deletion cassettes, the plasmid was digested with EciI.

*N. glabratus* deletion strains for *ERG3* and *ERG11* were generated in the ATCC2001 (CBS138) wild-type background. Overnight cultures in YPD medium were grown at 37 °C in a shaking incubator, followed by dilution to OD_600_ 0.4 in 50 mL YPD. Cells were incubated for three hours at 37 °C with shaking, harvested at 3500 rpm for 5 minutes, and washed twice with MilliQ water. The pellet was resuspended in 10 mL of 100 mM LiAc prepared in TE buffer (1x) and incubated at 37 °C for 30 minutes with shaking. DTT (1 M, 250 µL) was added, and cells were incubated for an additional hour. Ice-cold MilliQ water (40 mL) was added and cells were harvested (3500 rpm, 5 minutes, 4 °C), then washed once in ice-cold MilliQ water and once in 5 mL ice-cold sorbitol (1 M). The final pellet was resuspended in 500 µL ice-cold sorbitol (1 M). For electroporation, 1 µg of linearized deletion cassette was mixed with 40 µL of the prepared cells and transferred into a 2-mm gap electroporation cuvette. Cells were pulsed (1.5 kV, 200 Ω, 25 µF) and recovered in 2 mL YPD at 37 °C with shaking for three hours. Cells were spun down (4500 rpm, 2 minutes), resuspended in 100 µL MilliQ water, and plated on YPD agar supplemented with 200 µg/mL nourseothricin. For *ERG11* deletions, plates were further supplemented with 35 µg/mL ergosterol and 0.0875% (v/v) Tween 80 and incubated anaerobically at 37 °C, following recommendations from Geber et al. (11). Correct integration of the deletion cassette was confirmed by PCR using primers listed in **Table S3** (Supplementary).

For *ERG3* deletion strains, marker recycling was achieved via heat-shock with the pLS10 plasmid to express flippase. Overnight cultures in YPD medium were grown at 37 °C in a shaking incubator, followed by dilution to OD_600_ 0.4 in 50 mL YPD. Cells were incubated for three hours at 37 °C with shaking, harvested at 3500 rpm for 5 minutes, and washed twice with MilliQ water and once in 1 ml 100 mM LiAc. The pellet was resuspended in 500 μl 100 mM LiAc. For the transformation reaction, 50 µL cells were mixed with 240 µL 50% (w/v) PEG, 36 µL 1 M LiAc, 25 µL boiled Salmon Sperm DNA (2 mg/mL), and 1 µg of pLS10 plasmid in 50 µL MilliQ water. The mixture was incubated at 30 °C for 30 minutes in a shaking heat block, then heat-shocked at 42 °C for 22 minutes. Cells were recovered in 2 mL YPD at 37 °C for three hours, plated on YPD agar containing 300 µg/mL hygromycin, and incubated at 37°C. Loss of the nourseothricin resistance marker was confirmed by PCR using primers in **Table S3** (Supplementary). To remove the pLS10 plasmid, cells were plated on non-selective YPD medium and verified by replating on YPD supplemented with 300 µg/ml hygromycin.

### Drug and stress susceptibility testing

Antifungal susceptibility testing was done by the EUCAST reference method (15), Sensititer YeastOne 10 and ETEST® (bioMérieux) methods, following manufacturer’s instructions. An adaptation of the EUCAST method was used for drug and stress broth dilution assays (BDA). Briefly, a twofold dilution range of drug was prepared in a total volume of 200µL RPMI-MOPS (pH 7, 2% glucose, 1% DMSO) medium with approximately 200 cells (based on OD_600nm_ and serial dilution) in a round-bottom 96-well polystyrene microtiter plate (Greiner). The highest drug concentration was 16 µg/mL for all drugs except anidulafungin (max 8 µg/mL), micafungin (max 1µg/mL) and flucytosine (max 4µg/mL). All drugs except fluconazole were dissolved in 100% DMSO, while a final concentration of 1% DMSO was obtained in the final assay. Stressor dilution ranges were 10 - 0.0391 mM for hydrogen peroxide (H_2_O_2_; Sigma-Aldrich), calcofluor white (CFW; Fluorescent Brightener 28, Sigma-Aldrich) and Congo red (CR; Sigma-Aldrich), 2.5 - 0.0098 M for sodium chloride (NaCl, Sigma-Aldrich), 3.5 – 0.014mM for sodium dodecyl sulfate (SDS, Sigma-Aldrich) and 500 – 1.95 µM of bathophenanthrolinedisulfonate (BPS; Sigma-Aldrich) in the final concentration of the BDA. Plates were incubated at 37°C for 48 hours, and growth was assessed spectrophotometrically (OD_600_) using a Synergy™ H1 microplate reader (BioTek). Stress susceptibility was assessed using the same protocol.

The growth cut-off of all MIC values from BDA, was 50% growth compared to the drug-free control, while in ETEST® (bioMérieux), the MIC was indicated by the boundary of the growth inhibition zone on the strip, following manufacturers guidelines.

### Checkerboard assays

For checkerboard assays, the BDA protocol described above was used but two drugs were serially diluted across two gradients (one vertical for NIT, and one horizontal for AMB, 5FC or FLU). To evaluate drug interactive effects, the fractional inhibitory concentration index (FICI) was calculated and indicated in the right upper corner. *FICI* = *FIC*_*drug*1_ + *FIC*_*drug*2_ = 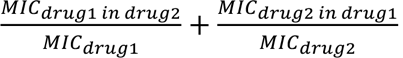, with MIC_drugA in drug B_ being the MIC of drug A in the presence of the highest concentration of drug B where growth is still visible (>50% compared to control without drug). FICI ≤ 0.5 indicates synergy, 0.5< FICI <1 indicates an additive effect, 1≤ FICI < 4 indicates indifference and a FICI > 4 indicates antagonism (33). For checkerboard assays in which the MIC of one of the two drugs could not be determined within the tested range, no FICI could be calculated.

### Growth analysis

Cultures were diluted in 200 µL RPMI-MOPS (with 0.2% or 2% glucose) or YPD (2% glucose) to a final cell concentration of 10^6^ cells per well. Growth was monitored at 37°C using spectrophotometric analysis at an optical density of 600 nm (OD_600_) with a Multiskan GO automated plate reader (Thermo Scientific) in flat-bottom 96-well microplates (Greiner) with intermittent (10 min. interval) pulsed (1 min medium strength shaking) shaking and 30-minute interval OD_600_ measurements. Growth curves were generated based on two replicate measurements per strain.

### Membrane sterol analysis

Sterols were extracted and analyzed based on Morio *et al.* (48) with some modifications. Stationary phase cultures were obtained by growing cells in 5 ml RPMI-MOPS (pH7, 2% glucose) medium in a shaking incubator at 37°C for 48 hours. Cells were harvested by centrifugation, washed twice with MilliQ H_2_O, and a pellet of 20 mg of cells was stored at −80°C. The pellet was resuspended (vortexing for 1 min.) in 300 µl saponification medium (12.5g KOH in 18ml MilliQ H_2_O diluted to 50ml with 98% ethanol), transferred to a capped glass vial and incubated for 1h in a shaking water bath at 80°C. Sterols were extracted by adding 100 µl MilliQ H_2_O and 400 µl hexane, including 1 µl of 5 mg/ml 5-α-cholestane as internal standard (Sigma, 47124), followed by vortexing for 3 min., 20 min. phase separation and careful collection of 350 µl of the top (hexane) layer. A second extraction fraction was collected by adding 600 µl hexane, vortexing for 3 min., 20 min. phase separation and collection of 550 µl of the top (hexane) layer. The two collected hexane fractions were combined and dried using vacuum centrifugation (Automatic Environmental SpeedVac® System AES2010) for 30 min at room temperature. Sterol extracts were re-dissolved in 60 µl hexane and derivatized by adding 10 µl of a silylating mixture (Sigma, 85432) short vortexing and incubation at room temperature for 1 h. Derivatized extracts were shortly centrifuged to precipitate potential debris and 50 µl of the extract was transferred to a smaller insert glass tube for GC-MS analysis.

The samples were analyzed using a Thermo Scientific gas chromatography-mass spectrometer [Trace 1300 - ISQ QD equipped with a TriPlus RSH autosampler and a Restek Rxi-5ms capillary GC column (30 m x 0.25mmID)]. Helium was used as carrier gas with a flow rate of 1.4 ml/min. Injection was carried out at 250°C in split mode after 1 min and with a ratio of 1:10. The temperature was first held at 50°C for 1 min and then allowed to rise to 260°C at a rate of 50°C/min, followed by a second ramp of 2°C/min until 325°C was reached; that temperature was maintained for 3 min. The mass detector was operated in scan mode (50 to 600 atomic mass units), using electron impact ionization (70 eV). The temperatures of the MS transfer line and detector were 325°C and 250°C, respectively. Sterols were identified by their retention time relative to the internal standard (cholestane) and specific mass spectrometric patterns using Chromeleon™ 7 (Thermo Scientific). The spectra were matched to GC-MS libraries described in Müller *et al.*(49) and NIST/EPA/NIH version 2. Analysis was performed by integration over the base ion of each sterol. Abundance was calculated relative to the internal standard, comparing the relative peak areas of the compounds.

Sterol extraction and analysis of each strain was performed in duplicate (technical repeats) on individually cultured strains.

### Confocal microscopy

Images shown in **Figure S2** (Supplementary) were taken by confocal microscopy. Cells grown on YPD agar were resuspended in RPMI-1640 MOPS (pH7) medium with 2% w/v glucose and samples were collected at two time points: immediately after inoculation (time 0 h) and after 24 hours of incubation at 37°C in a shaking incubator. Cells were harvested by centrifugation at 3,500 × g for 5 minutes and subsequent washing with phosphate-buffered saline (PBS). The pellets were resuspended in PBS containing 25 ng/mL calcofluor white (CFW; Sigma Aldrich) and 1 µL/mL LIVE/DEAD™ Fixable Olive stain (Thermo Fisher Scientific). The suspension was incubated for 30 minutes in darkness at room temperature and subsequently washed with PBS. Cells were visualized on glass slides using an Olympus Fluoview FV1000 inverted epi-fluorescence microscope equipped with a 60X (NA = 1.34, UPLSAPO) objective lens. Fluorescence excitation for CFW was performed using 405 nm light, while the LIVE/DEAD™ stain was excited using 488 nm light from an argon laser.

### Declaration of generative AI and AI-assisted technologies in the writing process

During the preparation of this work the author(s) used OpenAI GPT-o1 in order to aid in rephrasing, summarizing and proof-reading original written text. After using this tool/service, the author(s) reviewed and edited the content as needed and take(s) full responsibility for the content of the publication.

## Acknowledgements

This work was supported by the Fund for Scientific Research Flanders (FWO) under the framework of the JPIAMR – Joint Programming Initiative on Antimicrobial Resistance fund (project CycleDrug) and by a C3 grant from the Industrial Research Fund of KU Leuven (C3/22/007) granted to P.V.D. H.C. was supported by a post-doctoral fellowship granted by KU Leuven Internal Funds (PDMT2/23/032). V.B received funding from the European Union’s Horizon 2020 research and innovation program under the Marie Skłodowska-Curie grant agreement No 945352. J.V. and D.S., were supported by FWO PhD fellowships 11L0423N and 11J8122N, respectively. The T.G group acknowledges support from the Spanish Ministry of Science and Innovation (grant numbers PID2021-126067NB-I00, CPP2021-008552, PCI2022-135066-2, and PDC2022-133266-I00), cofounded by ERDF “A way of making Europe”, as well as support from the Catalan Research Agency (AGAUR) (grant number SGR01551); “La Caixa” foundation (grant number LCF/PR/HR21/00737), and Instituto de Salud Carlos III (IMPACT grant IMP/00019 and CIBERINFEC CB21/13/00061-ISCIII-SGEFI/ERDF). We would like to thank Dr. Caroline Mattelaer and the pathology department of ZAS for the histology images.

## Author contributions

H.C.: conceptualisation, study design, investigation, methodology, formal analysis, data curation, funding acquisition, visualisation and writing original draft. V.B., J.V.: investigation and methodology. C.L.R., J.P.H.A., R.V., D.S., G.V., L.V., B.B.X.: investigation. T.G., K.L., R.N., P.V.D.: supervision and funding acquisition. All authors edited and/or approved the manuscript.

## Supplementary

**Figure S1:**
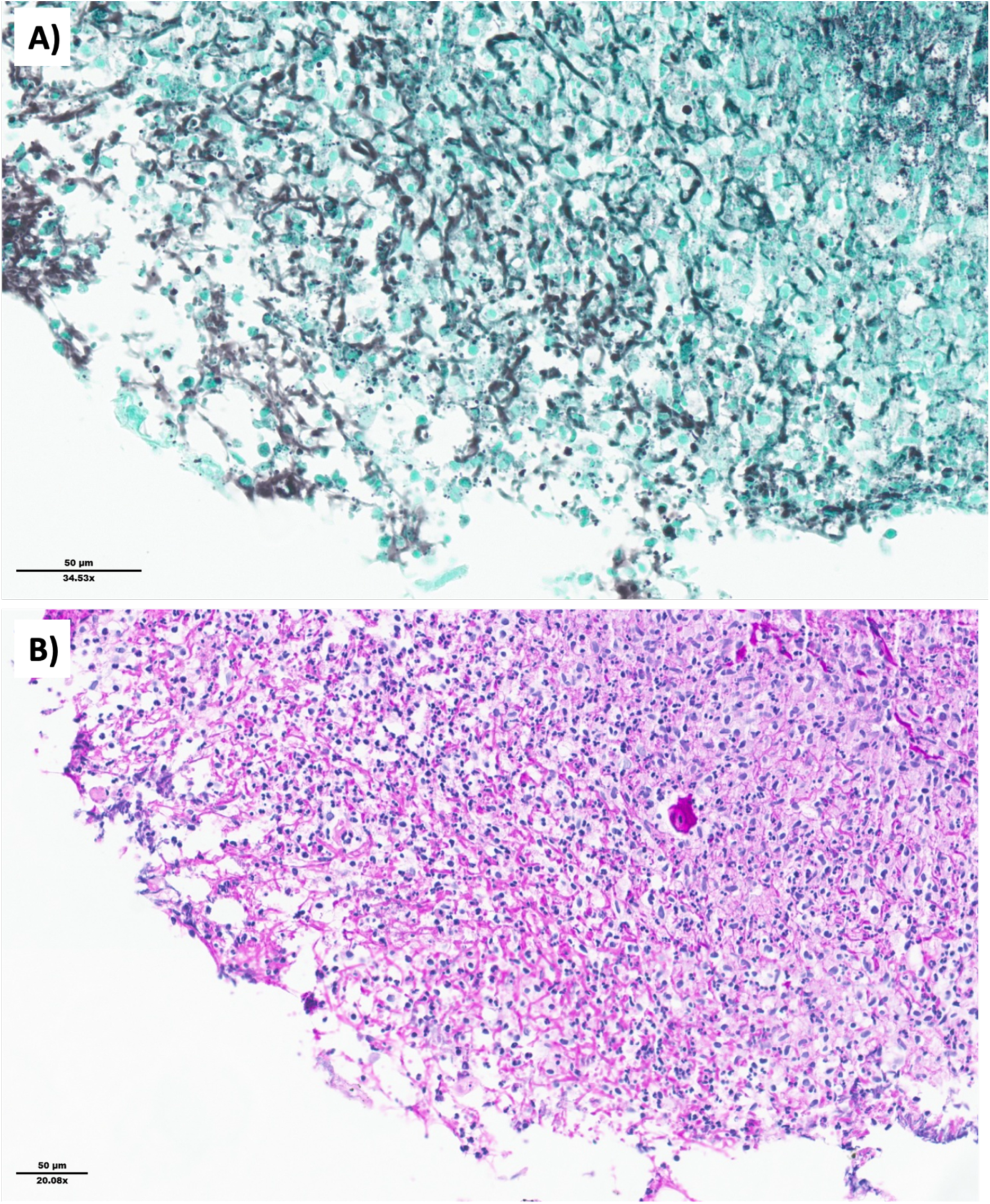
Formalin-fixed paraffin embedded sections of fibrinpurulent material from TURP show fungal structures in a Grocott-Gomori methenamine silver (GMS) staining (A) and inflammatory infiltration of immune cells in a Periodic acid–Schiff (PAS) amylase stain (B).

**Figure S2:**
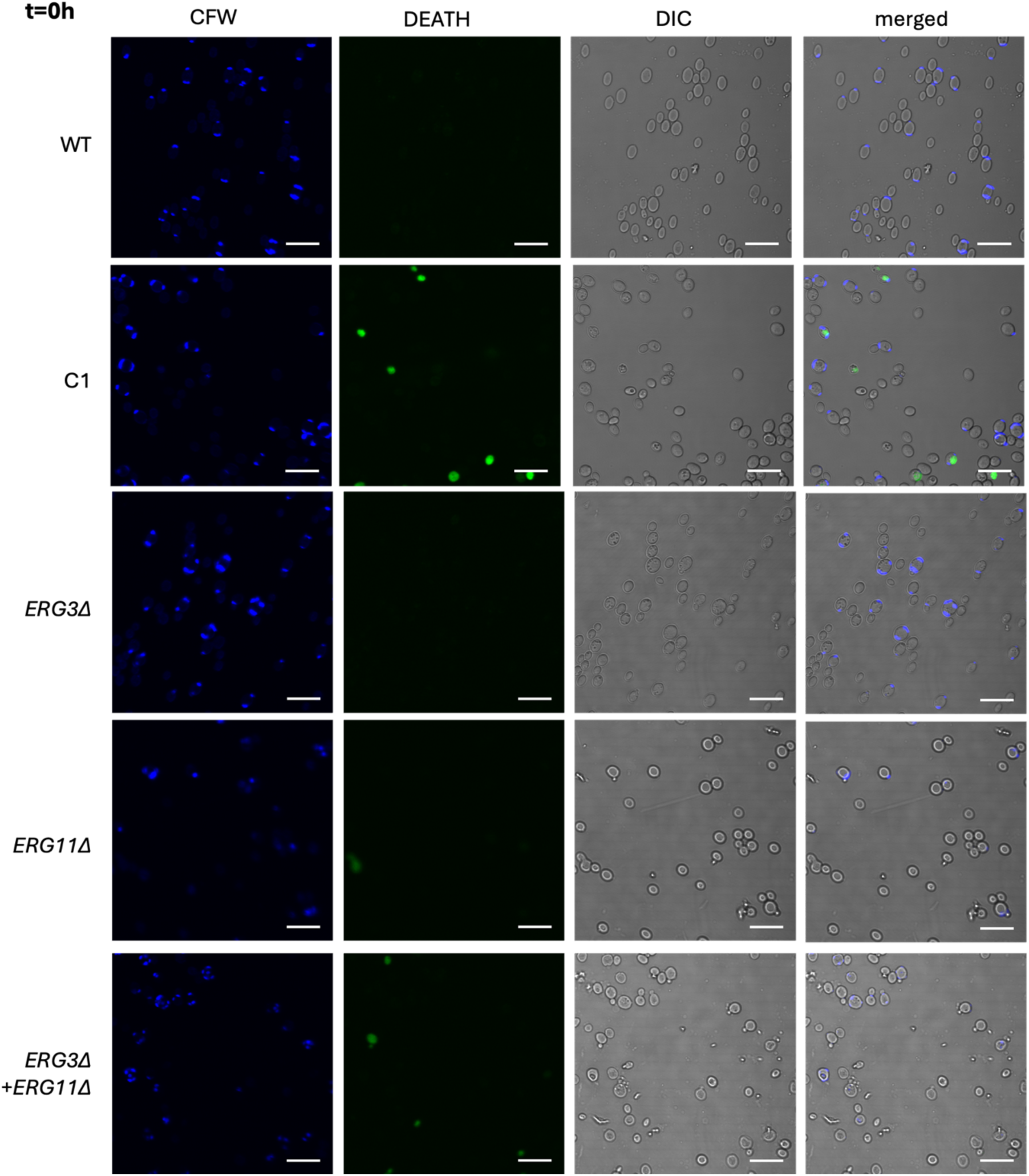
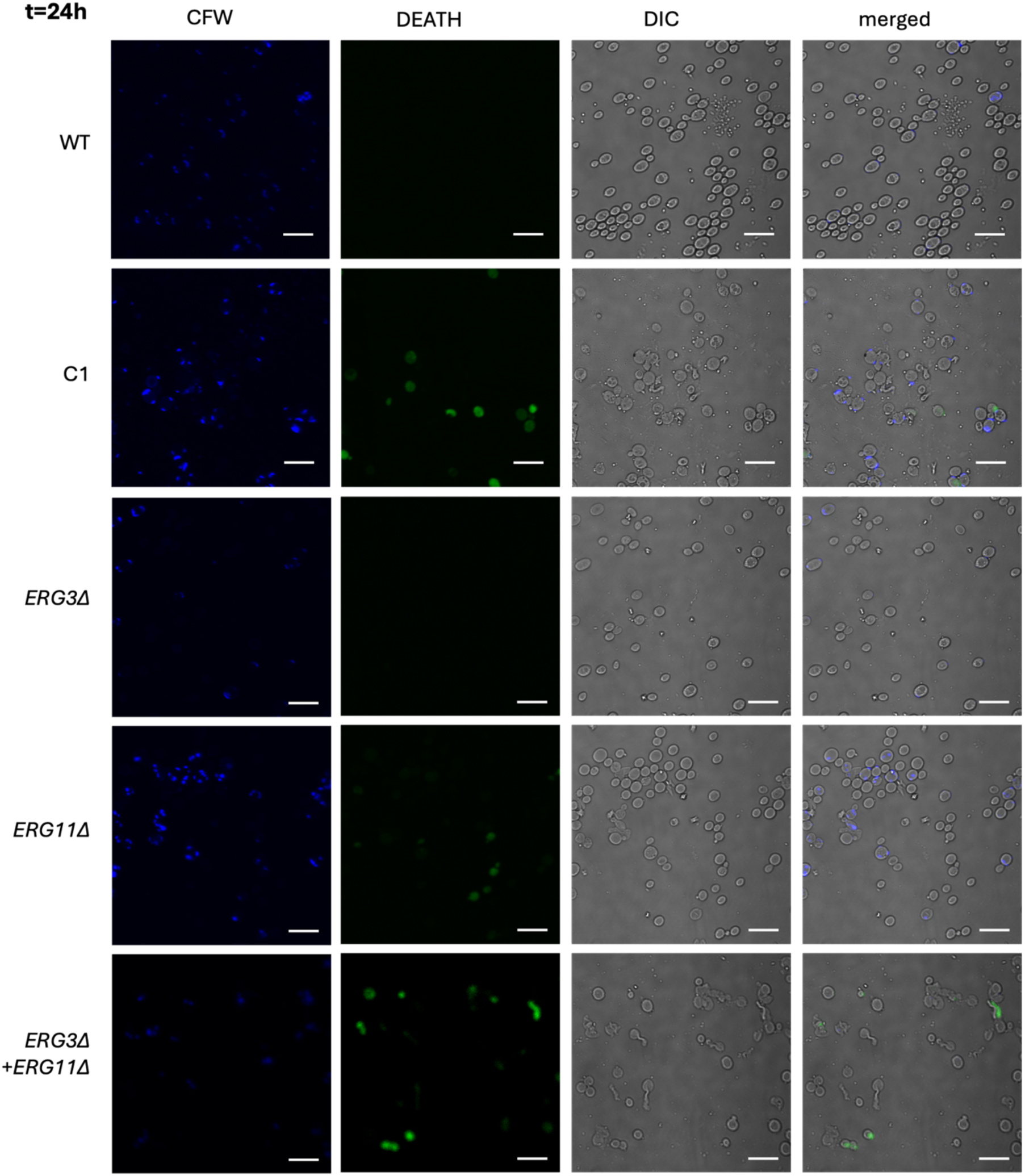
Differential interference contrast (DIC) confocal fluorescence microscopy of cells stained by calcofluor white (CFW) and LIVE/DEAD™ Fixable Olive stain (Thermo Fisher Scientific) after 0h and 24h of incubation in RPMI-MOPS 2% glucose at 37°C. CFW stains chitin and stains all fungal cells fluorescent blue. DEATH indicates the LIVE/DEAD stain, in which dead cells are fluorescent green.

**Figure S3:**
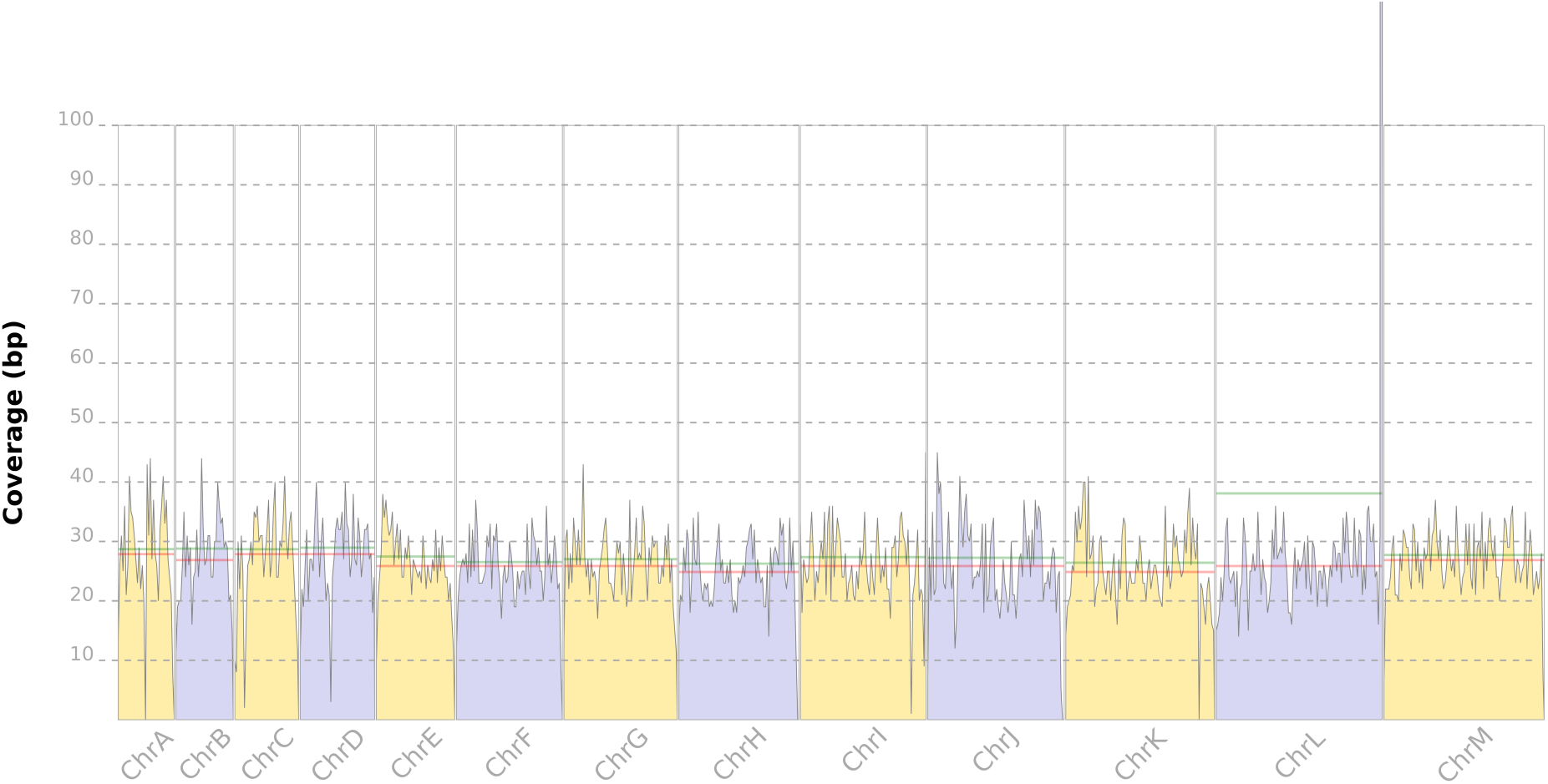
Genome-wide coverage plot of isolate C. Coverage depth (y-axis) is plotted against chromosomal positions (x-axis), with individual chromosomes labelled for clarity. The green and red lines represent the average and median coverage depth, respectively. The peak in coverage in ChrL corresponds to the cluster of rRNA genes, such as RDN25-1 (25S rRNA), RDN58-1 (5.8S rRNA), RDN18-1 (18S rRNA), and RDN5-1 (5S rRNA).

**Table S1:**
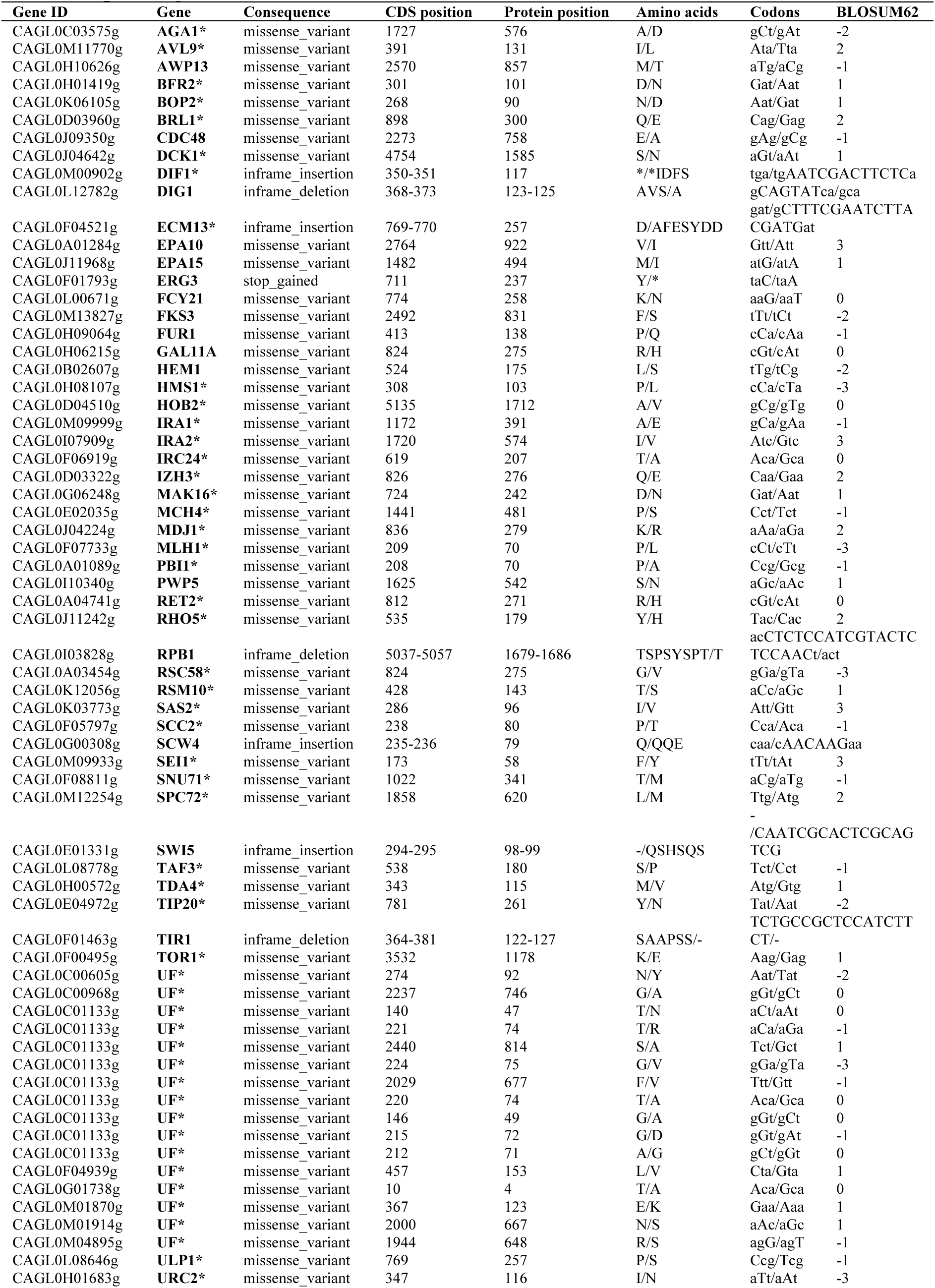
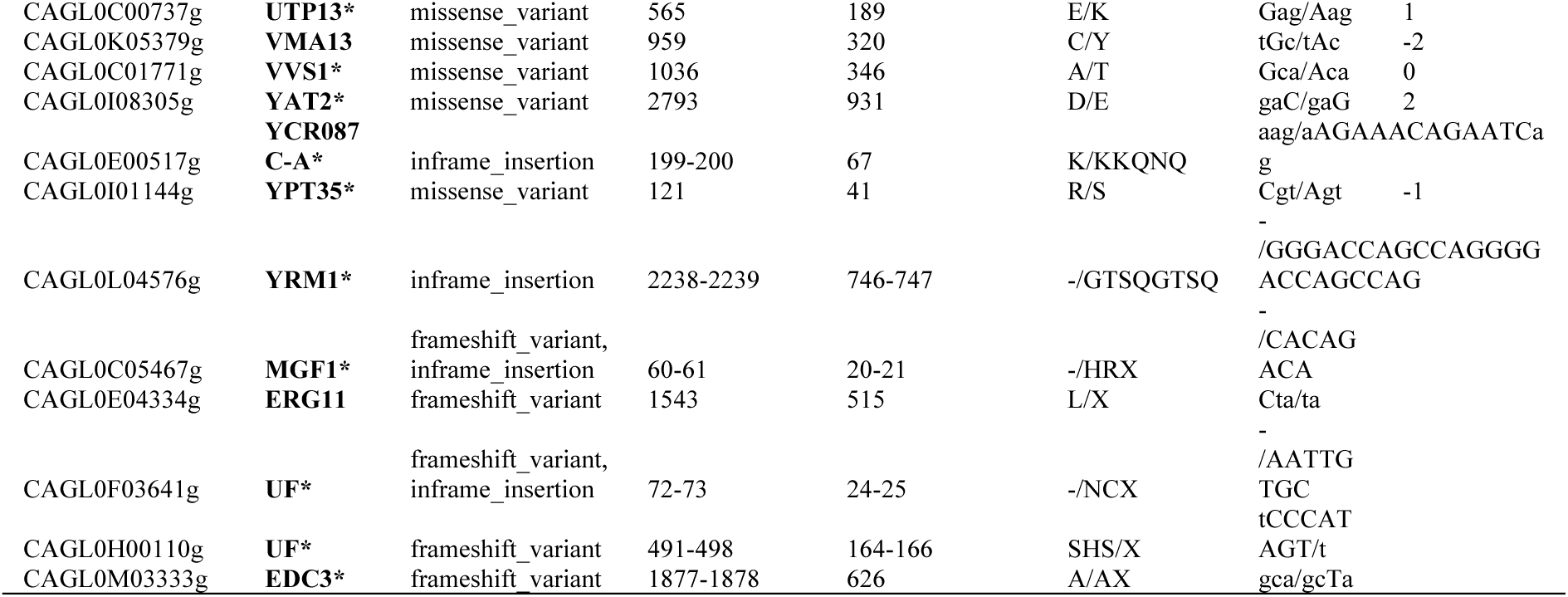
List of all protein-altering variants detected in the clinical isolate C. Given are gene IDs from the *N. glabratus* reference genome (ATCC2001/CBS6318), gene aliases or orthologous gene names in *S. cerevisiae* [marked with an asterisk (GENE*)], the mutation’s functional consequence, CDS and protein positions, specific amino acid and codon changes, and corresponding BLOSUM62 scores for amino acid substitutions.

**Table S2:**
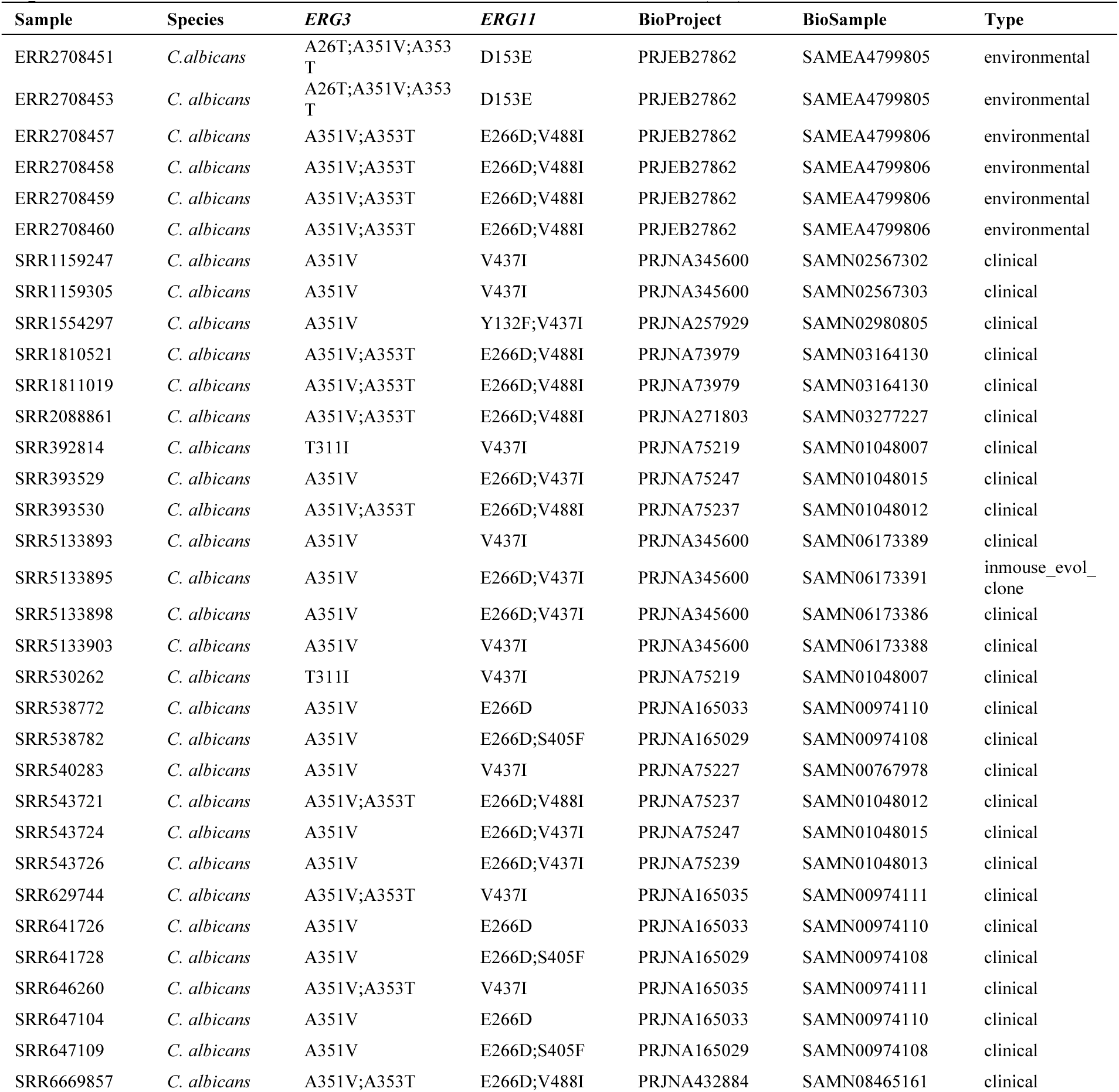
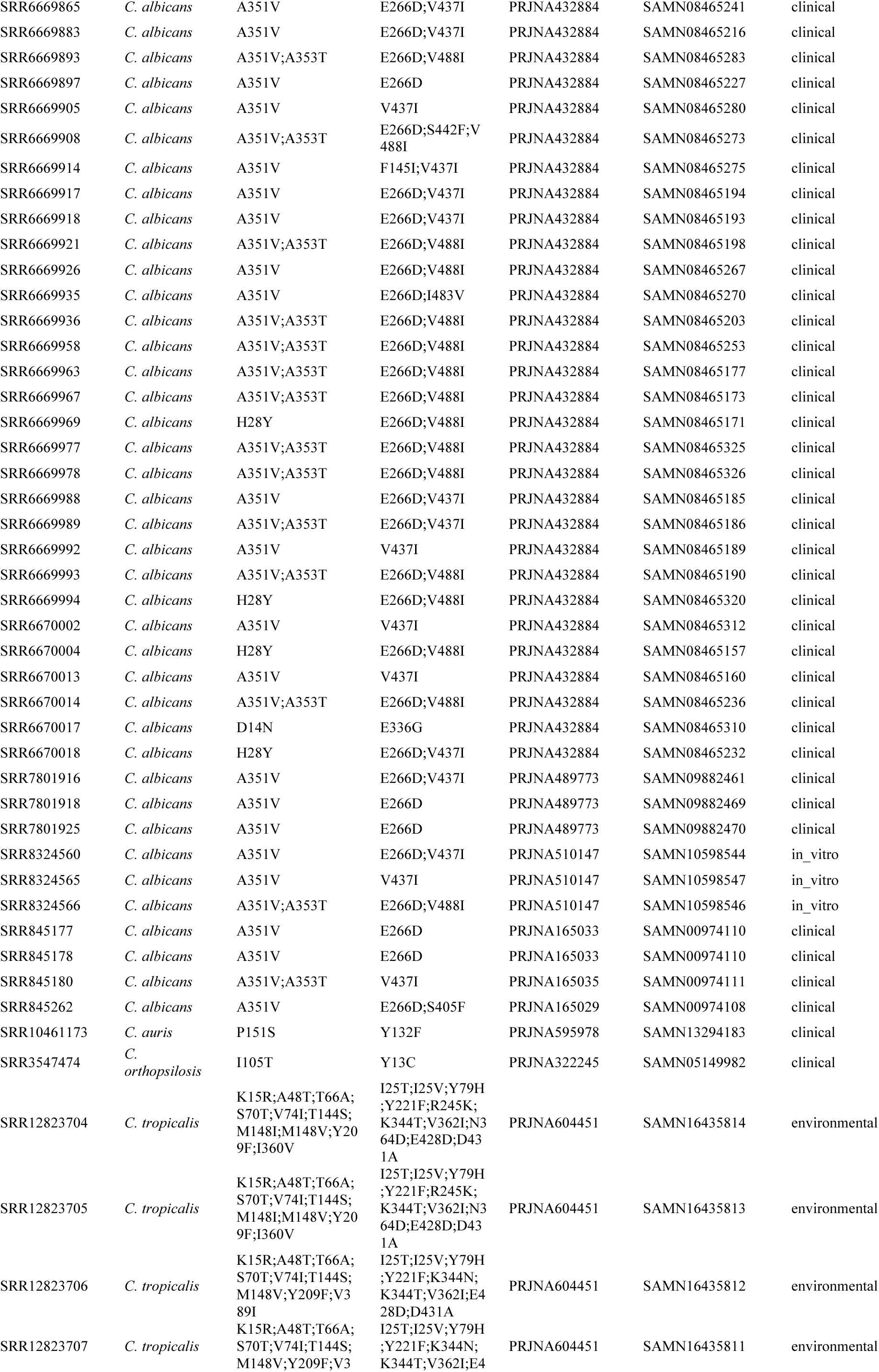
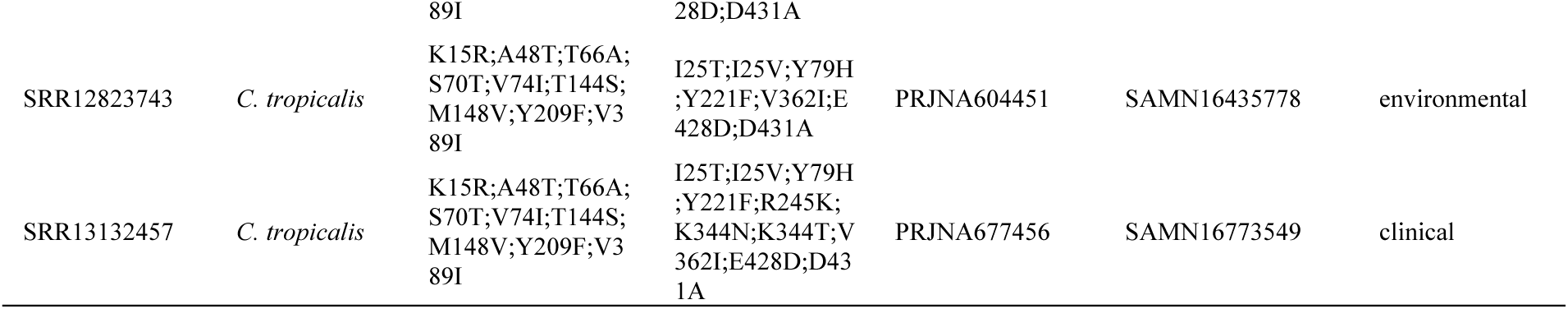
List of all samples containing combined *ERG3* and *ERG11* variants with frequency < 0.2. The table includes the SRR ID for each sample, species name, detected variants in ERG3 and ERG11, and corresponding BioProject and BioSample IDs. Additionally, the isolate type <u>is</u> provided as referenced in Schikora-Tamarit & Gabaldón (16).

**Table S3:**
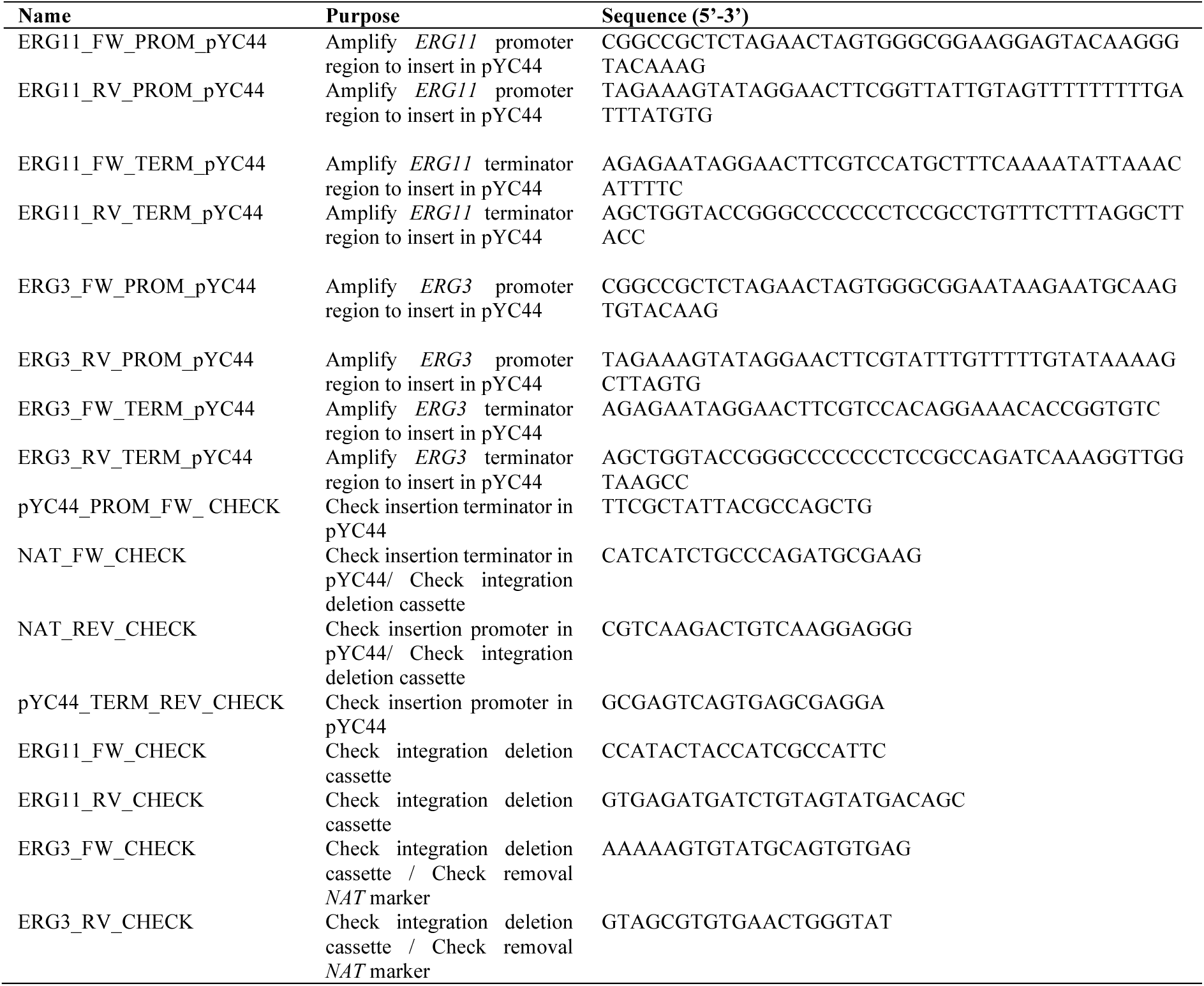
oligonucleotides used in this study.

## References

1. Denning DW. Global incidence and mortality of severe fungal disease. The Lancet Infectious Diseases. 2024;24(7):E428–E38.

2. Sharma M, Chakrabarti A. Candidiasis and Other Emerging Yeasts. Current Fungal Infection Reports. 2023;17(1):15–24.

3. Beardsley J, Kim HY, Dao A, Kidd S, Alastruey-Izquierdo A, Sorrell TC, et al. *Candida glabrata (Nakaseomyces glabrata)*: A systematic review of clinical and microbiological data from 2011 to 2021 to inform the World Health Organization Fungal Priority Pathogens List. Medical Mycology. 2024;62(6).

4. Behzadi P, Behzadi E, Ranjbar R. Urinary tract infections and *Candida albicans*. Central European Journal of Urology. 2015;68(1):96–101.

5. Sobel JD, Bradshaw SK, Lipka CJ, Kartsonis NA. Caspofungin in the treatment of symptomatic candiduria. Clinical Infectious Diseases. 2007;44(5):e46–9.

6. Pappas PG, Kauffman CA, Andes DR, Clancy CJ, Marr KA, Ostrosky-Zeichner L, et al. Clinical Practice Guideline for the Management of Candidiasis: 2016 Update by the Infectious Diseases Society of America. Clinical Infectious Diseases. 2015;62(4):e1–e50.

7. Carolus H, Pierson S, Munoz JF, Subotic A, Cruz RB, Cuomo CA, et al. Genome-Wide Analysis of Experimentally Evolved *Candida auris* Reveals Multiple Novel Mechanisms of Multidrug Resistance. mBio. 2021;12(2):e03333–20.

8. Vincent BM, Lancaster AK, Scherz-Shouval R, Whitesell L, Lindquist S. Fitness Trade-offs Restrict the Evolution of Resistance to Amphotericin B. PLOS Biology. 2013;11(10):e1001692.

9. Carolus H, Sofras D, Boccarella G, Septhon-Clark P, Biriukov V, Cauldron NC, et al. Acquired amphotericin B resistance leads to fitness trade-offs that can be mitigated by compensatory evolution in *Candida auris*. Nature Microbiology. 2024;9:3304–20

10. Carolus H, Pierson S, Lagrou K, Van Dijck P. Amphotericin B and Other Polyenes-Discovery, Clinical Use, Mode of Action and Drug Resistance. Journal of Fungi. 2020;6(4):321.

11. Geber A, Hitchcock CA, Swartz JE, Pullen FS, Marsden KE, Kwon-Chung KJ, et al. Deletion of the *Candida glabrata ERG3* and *ERG11* genes: effect on cell viability, cell growth, sterol composition, and antifungal susceptibility. Antimicrobial Agents and Chemotherapy. 1995;39(12):2708–17.

12. Sanglard D, Ischer F, Parkinson T, Falconer D, Bille J. *Candida albicans* mutations in the ergosterol biosynthetic pathway and resistance to several antifungal agents. Antimicrobial Agents and Chemotherapy. 2003;47(8):2404–12.

13. Eddouzi J, Parker JE, Vale-Silva LA, Coste A, Ischer F, Kelly S, et al. Molecular mechanisms of drug resistance in clinical *Candida* species isolated from Tunisian hospitals. Antimicrobial Agents and Chemotherapy. 2013;57(7):3182–93.

14. CLSI. CLSI M57S: Epidemiological Cutoff Values for Antifungal Susceptibility Testing, 4th Edition 2022 [Available from: https://clsi.org/standards/products/microbiology/documents/m57s/.

15. EUCAST. Breakpoint tables for interpretation of MICs for antifungal agents Version 11.0, valid from 2024-12-02 2024 [Available from: https://www.eucast.org/fileadmin/src/media/PDFs/EUCAST_files/AFST/Clinical_breakpoints/AFST_BP_v11.0.pdf.

16. Schikora-Tamarit MA, Gabaldón T. Recent gene selection and drug resistance underscore clinical adaptation across *Candida* species. Nature Microbiology. 2024;9(1):284–307.

17. Carrete L, Ksiezopolska E, Pegueroles C, Gomez-Molero E, Saus E, Iraola-Guzman S, et al. Patterns of Genomic Variation in the Opportunistic Pathogen *Candida glabrata* Suggest the Existence of Mating and a Secondary Association with Humans. Current Biology. 2022;32(14):3219.

18. Hecht M, Bromberg Y, Rost B. Better prediction of functional effects for sequence variants. BMC Genomics. 2015;16.

19. Henikoff S, Henikoff JG. Amino acid substitution matrices from protein blocks. Proceedings of the National Academy of Sciences. 1992;89(22):10915–9.

20. Cline MS, Karchin R. Using bioinformatics to predict the functional impact of SNVs. Bioinformatics. 2011;27(4):441–8.

21. Delma FZ, Al-Hatmi AMS, Bruggemann RJM, Melchers WJG, de Hoog S, Verweij PE, et al. Molecular Mechanisms of 5-Fluorocytosine Resistance in Yeasts and Filamentous Fungi. Journal of Fungi. 2021;7(11).

22. Paluszynski JP, Klassen R, Rohe M, Meinhardt F. Various cytosine/adenine permease homologues are involved in the toxicity of 5-fluorocytosine in *Saccharomyces cerevisiae*. Yeast. 2006;23(9):707–15.

23. Bédard C, Pageau A, Fijarczyk A, Mendoza-Salido D, Alcañiz AJ, Després PC, et al. FungAMR: A comprehensive portrait of antimicrobial resistance mutations in fungi. bioRxiv (preprint). 2024:2024.10.07.617009.

24. Fuchs F, Aldejohann AM, Hoffmann AM, Walther G, Kurzai O, Hamprecht AG. *In Vitro* Activity of Nitroxoline in Antifungal-Resistant *Candida* Species Isolated from the Urinary Tract. Antimicrobial Agents and Chemotherapy. 2022;66(6):e0226521.

25. Cherdtrakulkiat R, Boonpangrak S, Sinthupoom N, Prachayasittikul S, Ruchirawat S, Prachayasittikul V. Derivatives (halogen, nitro and amino) of 8-hydroxyquinoline with highly potent antimicrobial and antioxidant activities. Biochemistry and Biophysics Reports. 2016;6:135–41.

26. Fuchs F, Hof H, Hofmann S, Kurzai O, Meis JF, Hamprecht A. Antifungal activity of nitroxoline against *Candida auris* isolates. Clinical Microbiology and Infection. 2021;27(11):1697–10.

27. Carolus H, Sofras D, Boccarella G, Jacobs S, Biriukov V, Goossens L, et al. Collateral sensitivity counteracts the evolution of antifungal drug resistance in *Candida auris*. Nature Microbiology. 2024;9:2954–69.

28. Song Yn, Xu H, Chen W, Zhan P, Liu X. 8-Hydroxyquinoline: a privileged structure with a broad-ranging pharmacological potential. MedChemComm. 2015;6:61–74.

29. de Chaves MA, da Costa BS, de Souza JA, Batista MA, de Andrade SF, Hage-Melim LIdS, et al. In silico and in vitro analysis of the mechanisms of action of nitroxoline against some medically important opportunistic fungi. Journal of Medical Mycology. 2023;33(3):101411.

30. Cacace E, Tietgen M, Steinhauer M, Mateus A, Schultze TG, Eckermann M, et al. Uncovering nitroxoline activity spectrum, mode of action and resistance across Gram-negative bacteria. bioRxiv (preprint). 2024:2024.06.04.597298.

31. Subissi A, Monti D, Togni G, Mailland F. Ciclopirox. Drugs. 2010;70(16):2133–52.

32. Cheng Y-S, Roma JS, Shen M, Fernandes CM, Tsang PS, Forbes HE, et al. Identification of Antifungal Compounds against Multidrug-Resistant *Candida auris* Utilizing a High-Throughput Drug-Repurposing Screen. Antimicrobial Agents and Chemotherapy. 2021;65(4).

33. Doern CD. When Does 2 Plus 2 Equal 5? A Review of Antimicrobial Synergy Testing. Journal of Clinical Microbiology. 2014;52(12):4124–8.

34. Whaley SG, Berkow EL, Rybak JM, Nishimoto AT, Barker KS, Rogers PD. Azole Antifungal Resistance in *Candida albicans* and Emerging Non-*albicans Candida* Species. Frontiers in Microbiology. 2017;7.

35. Kelly SL, Lamb DC, Kelly DE, Manning NJ, Loeffler J, Hebart H, et al. Resistance to fluconazole and cross-resistance to amphotericin B in *Candida albicans* from AIDS patients caused by defective sterol Δ5,6-desaturation. FEBS Letters. 1997;400(1):80–2.

36. Gregor JB, Gutierrez-Schultz VA, Hoda S, Baker KM, Saha D, Burghaze MG, et al. An expanded toolkit of drug resistance cassettes for *Candida glabrata*, Candida auris, and Candida albicans leads to new insights into the ergosterol pathway. mSphere. 2023;8(6):e00311–23.

37. Hill JA, Cowen LE. Using combination therapy to thwart drug resistance. Future Microbiology. 2015;10(11):1719–26.

38. Torella JP, Chait R, Kishony R. Optimal Drug Synergy in Antimicrobial Treatments. PLOS Computational Biology. 2010;6(6):e1000796.

39. Chait R, Craney A, Kishony R. Antibiotic interactions that select against resistance. Nature. 2007;446(7136):668-71.

40. Munck C, Gumpert HK, Wallin AI, Wang HH, Sommer MO. Prediction of resistance development against drug combinations by collateral responses to component drugs. Science Translational Medicine. 2014;6(262):262ra156.

41. Rodriguez de Evgrafov M, Gumpert H, Munck C, Thomsen TT, Sommer MOA. Collateral Resistance and Sensitivity Modulate Evolution of High-Level Resistance to Drug Combination Treatment in *Staphylococcus aureus*. Molecular Biology and Evolution. 2015;32(5):1175–85.

42. Schikora-Tamarit MA, Gabaldon T. PerSVade: personalized structural variant detection in any species of interest. Genome Biology. 2022;23(1):175.

43. Andrews S. FastQC: A Quality Control Tool for High Throughput Sequence Data 2010 [Available from: http://www.bioinformatics.babraham.ac.uk/projects/fastqc/.

44. Bolger AM, Lohse M, Usadel B. Trimmomatic: a flexible trimmer for Illumina sequence data. Bioinformatics. 2014;30(15):2114–20.

45. Danecek P, Bonfield JK, Liddle J, Marshall J, Ohan V, Pollard MO, et al. Twelve years of SAMtools and BCFtools. Gigascience. 2021;10(2).

46. Poplin R, Ruano-Rubio V, DePristo MA, Fennell TJ, Carneiro MO, Auwera GAVd, et al. Scaling accurate genetic variant discovery to tens of thousands of samples. bioRxiv (preprint). 2018.

47. Garisson E, Marth G. Haplotype-based variant detection from short-read sequencing (preprint). arXiv (preprint). 2012.

48. Morio F, Pagniez F, Lacroix C, Miegeville M, Le Pape P. Amino acid substitutions in the *Candida albicans* sterol Δ5,6-desaturase (Erg3p) confer azole resistance: characterization of two novel mutants with impaired virulence. Journal of Antimicrobial Chemotherapy. 2012;67(9):2131–8.

49. Müller C, Binder U, Bracher F, Giera M. Antifungal drug testing by combining minimal inhibitory concentration testing with target identification by gas chromatography-mass spectrometry. Nature Protocols. 2017;12(5):947–63.

